# A scalable pipeline for SARS-CoV-2 replicon construction based on de-novo synthesis

**DOI:** 10.1101/2022.02.05.478644

**Authors:** Ahmet C. Berkyurek, Elian Lee, Omer Weissbrod, Roni Rasnic, Ilaria Falciatori, Siyuan Chen, Yaniv Erlich

## Abstract

Replicons are synthetic viral RNA molecules that recapitulate the self-replicating activities of the virus but are missing its infectivity potential. Here, we report on a scalable pipeline to generate a replicon of any SARS-CoV-2 strain using de-novo synthesis. Our pipeline relies only on publicly available sequencing data without requiring access to any material, simplifying logistical and bureaucratic issues of sample acquisition. In addition, our system retains the nucleotide sequence of most of the SARS-CoV-2 full genome and therefore better captures its underlying genomic and biological functions as compared to the popular pseudotypes or any replicon system published to-date. We utilized our system to synthesize a SARS-CoV-2 non-infectious version of the Beta strain. We then confirmed that the resulting RNA molecules are non-infectious and safe to handle in a BSL2/CL2 facility. Finally, we show that our replicon can be specifically inhibited by molnupiravir and RNAi treatments, demonstrating its utility for drug research and development.

## Introduction

The constant emergence of SARS-CoV-2 variants necessitates a system for rapid assessment and prototyping of pharmacological interventions. Sequencing data of new strains is published and shared at an internet speed, but obtaining a live virus sample for preclinical studies is a much slower process. Based on our experience, there are three logistical factors that typically hinder access to live virus samples. First, sourcing the new strains is not always straightforward. The process typically requires identifying a collaborator with a live strain sample and with the capacity and motivation to collaborate. Second, getting the virus often entails a material transfer agreement, which includes legal review and additional red tape, exacerbating the logistical complexity, and tends to incur lengthy delays. Finally, once the viral sample arrives, the experiments require access to one of the BSL-3 facilities, which are in high demand. Indeed, multiple studies have utilized pseudotyping as an alternative for live virus experiments ^1,2^. However, this approach can only model Spike mutations but not other types of changes in the viral genome.

Replicon systems are a safe alternative for live virus experiments^3^. These systems rely on deleting from the viral genome one or more genes obligatory for its assembly, while maintaining the replicon core intact. The modified viral genome is then introduced into the target cells as naked RNA, where it is capable of in-cell replication, but cannot exit the cells because it misses key proteins for virion release. This approach has been utilized in multiple positive strand RNA viruses, from alphavirus to human hepatitis C virus to SARS-CoV-2^3–9^. However, most of these efforts require the viral genome as their initial material for the replicon system, again introducing the same logistical issues of obtaining an initial sample ^6,8,9^. Other studies removed large parts of the subgenomic regions, precluding targeting these regions by therapeutic agents ^4,7,10^.

Here, we developed and validated a scalable pipeline for creating safe and effective replicons at will, and demonstrated its utility for SARS-CoV-2 research and drug development. By harnessing the power of massively parallel DNA synthesis, our pipeline is a fully *ab-initio* system, free from any legal or biological strings attached to a specific biological sample, allowing fast strain prototyping. The pipeline starts by bioinformatically screening all publicly available sequencing data to create a consensus sequence for a specific strain of interest. Next, it automatically introduces a small number of synthetic alterations, aimed to completely eliminate viral assembly using multiple layers of protection, while maximising retention of the integrity of the rest of the viral genome. In addition, we integrate a GFP-nLuc encoding sequence into the replicon, to enable quick and cost-effective tracing of its activity. To demonstrate the capabilities of this pipeline, we synthesized a SARS-CoV-2 Beta (B.1.351) replicon, verified its safety, and demonstrated its utility for therapeutic research and development by subjecting it to multiple pharmaceutical manipulations.

## Results

### Automatic pipeline to design a safe replicon system using public data

To design the replicon, we created a bioinformatics pipeline that harnesses the vast amount of publicly available SARS-CoV-2 sequencing data deposited by the international community. The process starts by specifying the lineage of the desired strain. Then, it quickly scans the SARS-CoV-2 genomes to find the required strains at the NCBI database, which we prioritised over other resources as there are no commercial restrictions for using the data. It then performs multiple sequence alignment to create a consensus sequence. Finally, it annotates the genome to identify each genetic element in the ancestral strain (**Figure 1A**).

**Figure 1:**
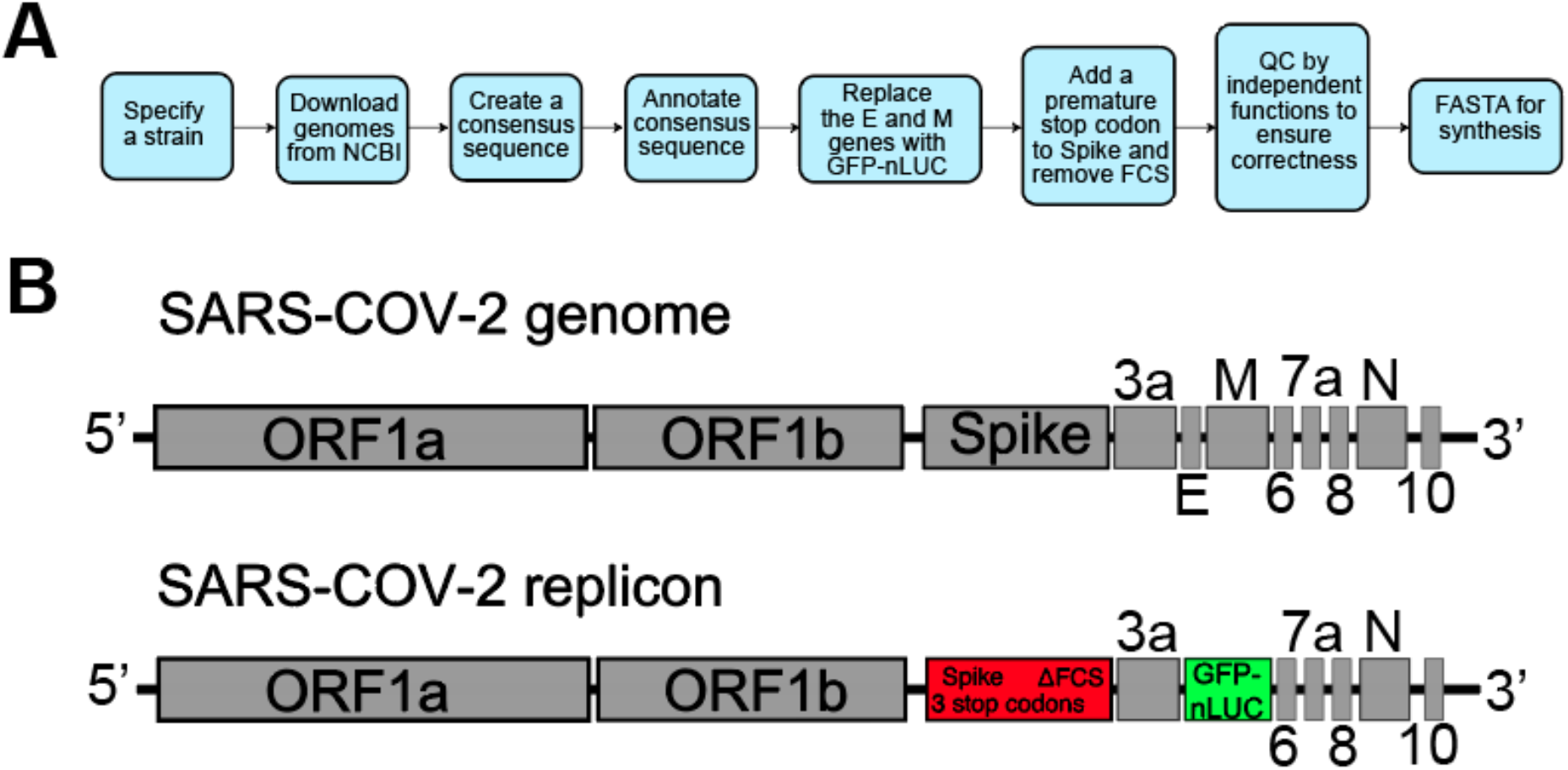
Establishing a safe SARS-CoV-2 replicon. **A.** Bioinformatics pipeline to design the SARS-CoV-2 replicon. **B.** Schematic illustration of the SARS-CoV-2 replicon against the sequence of the viral genome. E and M genes are deleted and replaced with a GFP-nLuc construct. The furin cleavage site (FCS) on the Spike gene was deleted and 3 early stop mutations were introduced.

Next, the algorithm designs assembly-prevention alterations by fully deleting the Membrane (M) and the Envelope (E) genes from the replicon genome. The M protein is required for virion production in coronaviruses^11^. It is the most abundant structural protein, found in virtually all branches of Coronavirinae^12^, and necessary for viral production^11^. A segment of its transmembrane domain is highly conserved, to the point that it is also found in torovirus, the closest evolutionary outer group to Coronavirinae^12^, which reflects its importance. The E gene, encoding for the virus’s envelope protein, is less conserved, but has been shown to be required for maintaining viral pathogenesis^12^. Studies of SARS-CoV have shown that deletion of the E gene results in a highly attenuated virus^11^ and that both the E and M genes are necessary for the formation of SARS-CoV virus-like particles^13^. Taken together, these multiple lines of evidence indicate that the M and E genes are essential for SARS-CoV-2 infectivity.

To further verify the role of the M and E genes, we analyzed the mutation profiles of over 26,000 SARS-CoV-2 genomes. We calculated the probability to observe a nonsense mutation in each gene adjusted to its length. Our analysis shows that the E and M genes are two of the top three structural proteins with the smallest number of nonsense mutations (**Supp. Table 1**). Other structural proteins, such as ORF8, ORF6, and N exhibited more than three fold the rate of loss of function mutations. As expected, the genetic pressure against deleterious effects in viable virions mirrors their importance to viral survival.

Instead of the E and the M genes, we introduced a GFP-nLuc-encoding sequence, which enables tracing the expression of subgenomic genes either using bioluminescence- or fluorescence-based experiments. The construct is driven by the E promoter and therefore the GFP-nLuc sequence behaves like a subgenomic RNA region.

In addition to deleting the M and E genes, our pipeline suppresses the expression of the Spike protein. To this end, our pipeline introduces a premature UAA stop codon instead of the 10th amino-acid in the sequence of Spike. Following the stop codon, our pipeline also introduces a frameshift mutation by adding a single “G” nucleotide after the premature stop codon. The result is a structure that is robustly resistant to viral escape mechanisms via deletion of any single A from the stop codon or even of the entire stop codon. Lastly, we also removed the Furin Cleavage Site (FCS) from the Spike region. Previous studies have shown that deletion of the FCS substantially attenuates virulence^14^, providing an additional safety feature in the extremely unlikely case of viral assembly. We kept the rest of the Spike region intact, aiming to enable testing therapies targeting its nucleic acid sequence and preserving as much as possible of the structural aspects of the RNA genome (**Figure 1B; Supplemental Data File 1**).

Based on all of the above, the safety committee of the University of Cambridge, which regulates our experiments, assigned a BSL2/CL2 level to the viral replicon.

### *De novo* construction of a safe SARS-CoV-2 replicon system

We used our bioinformatics pipeline to rapidly establish a replicon system in the lab. We selected B.1.351 as our strain of interest and constructed a consensus sequence. We then harnessed Twist Bioscience’s massively parallel DNA synthesis array to construct the replicon system, while adding a T7 site at the 5’ end to drive RNA transcription. Briefly, 1.6kb adaptor-on non-clonal gene fragments were synthesized by Twist to be used as building blocks for the full 30kb replicon with a turnaround time of 7 days. The building blocks were first assembled into 5kb segments, followed by another assembly into 15kb fragments *in vitro,* which took ~3 days. Over the span of the following week, the 2 final 15kb fragments were gel-purified before being cloned into a BAC vector and electroporated into BAC replicator cells. Clones were picked and verified by NGS as the accurate 30kb replicon construct, at which point they were glycerol banked. Overall, the process took about 3 weeks.

Next, we used T7 RNA polymerase to transcribe the replicon using an optimized protocol (**Figure 2A**). Initial attempts to transcribe the long DNA molecule have proven challenging and resulted in incomplete products and low yields (**Figure 2B**). We eventually found that overactivating the T7 RNA polymerase by HEPES buffer, and reducing the temperature down to 27°C, increased transcription processivity and enabled high fidelity production of full length transcripts. Using this protocol, we were able to repeatedly generate yields of 4ug/ul of replicon RNA in a total volume of 100ul (**Figure 2C**).

**Figure 2:**
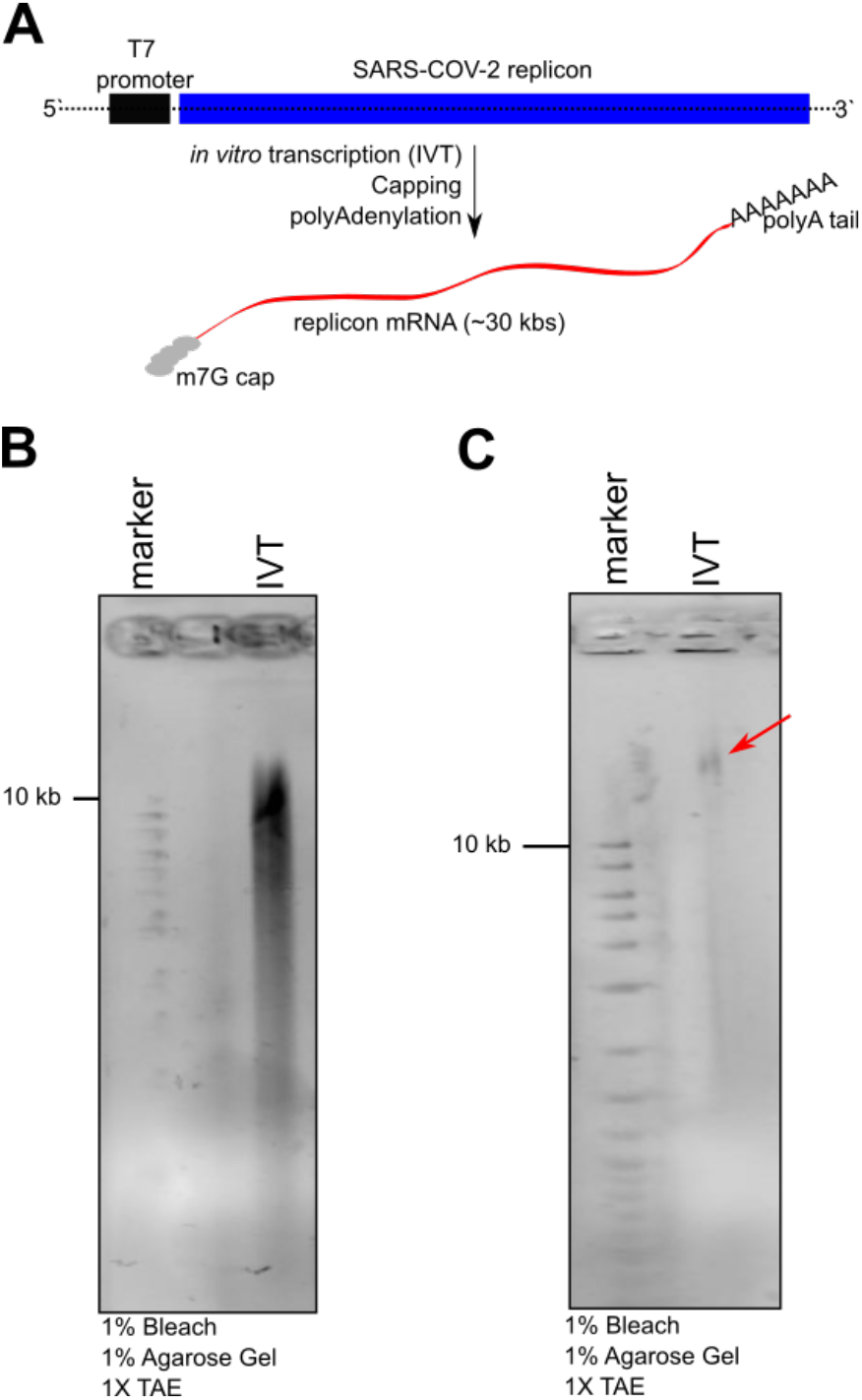
*in vitro* transcription (IVT) of the SARS-CoV-2 replicon: **A.** Illustration of the in *vitro* transcription for the linearised plasmid. **B.** Initial attempts produced incomplete products shorter than 30 kb. **C.** Addition of 50 mM HEPES (pH=7.5), 10 mM NaCl and reducing the temperature to 27°C give rise to the full product, at the expected size of 30 kb. (Red arrow indicates the intact *in vitro* transcribed replicon mRNA).

We confirmed the safety of the replicon products via multiple methods. First, we sequenced the DNA template of the replicon to confirm its design using Illumina sequencing. We identified a perfect match with the desired target sequence, did not identify any mutations in the replicon product as compared to its desired FASTA design, validated that the E and M genes were indeed removed, and confirmed that the Spike gene contained the two required mutations (**Figure 3A**). Second, we transfected the replicon in HEK293, BHK21, and VeroE6 cells and measured its expression after 24 hours by qPCR using the Berlin protocol that amplifies the RdRP and E gene levels^15^. Our results showed strong Ct values of RdRP (15 to 22), but no signal from the E gene, confirming the absence of activity from this gene (**Figure 3B**). Third, we conducted an infection experiment, which is the ultimate test for the safety of the replicon. In this experiment, we transfected VeroE6 cells with the replicon and challenged fresh VeroE6 cells with the supernatant of the transfected cells 48 hours post-transfection. We then tested the replicon level in both the transfected VeroE6 cells and the challenged ones using qPCR. We failed to detect any RdRP levels in the challenged cells whereas the transfected cells showed a Ct level of 12 (**Figure 3C**). This confirmed that our replicon product could not infect naive cells that are otherwise highly susceptible to viral infection.

**Figure 3:**
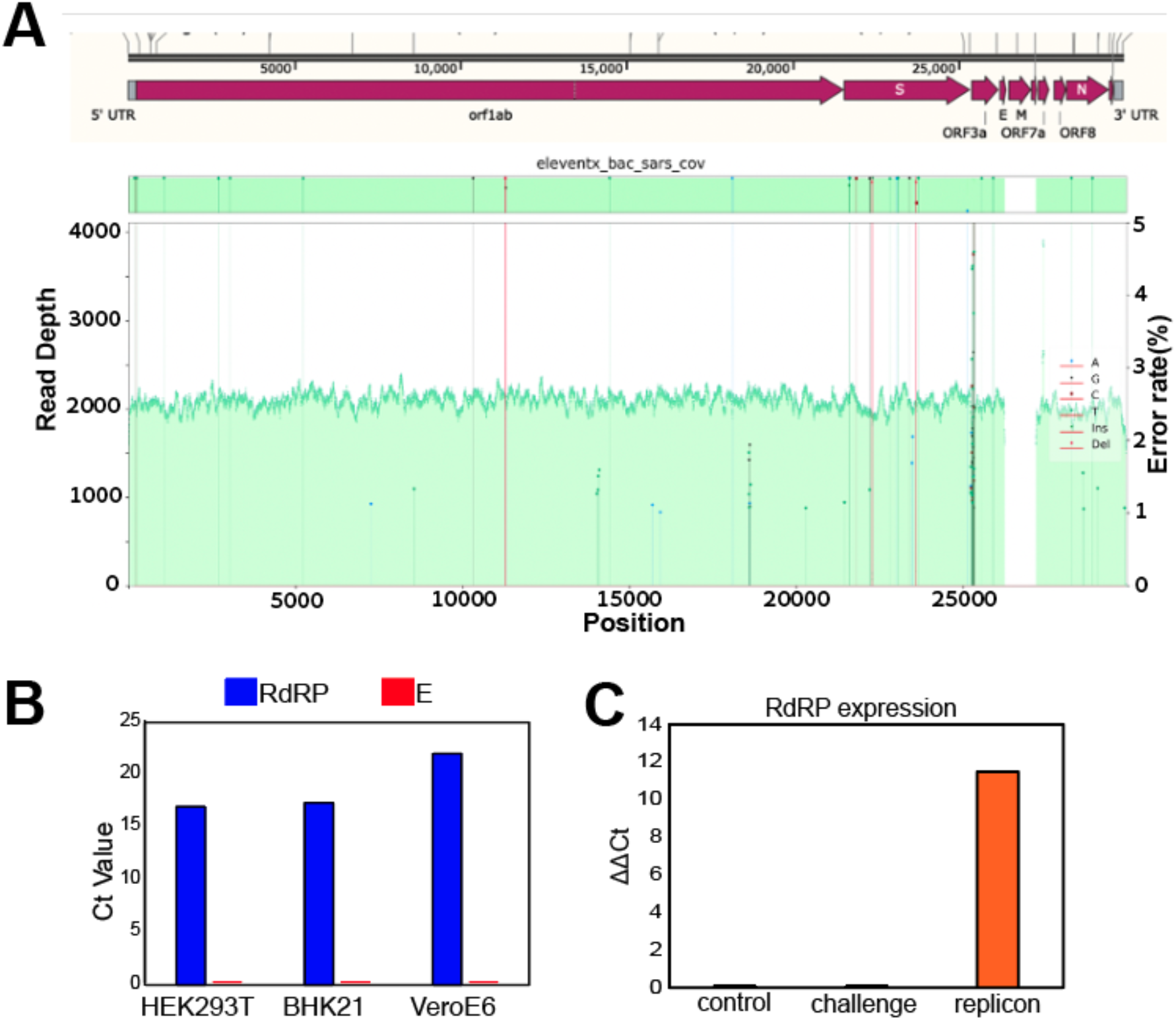
The SARS-CoV-2 replicon is non-infective and safe: **A.** DNA Sequencing of the replicon plasmid showed that the structural genes E and M were missing from our system. **B.** qPCR expression of the RdRP and E genes in replicon-transfected cells (HEK293T, BHK21 and VeroE6). Ct values for the E gene could not be detected. **C.** The supernatant from replicon transfected VeroE6 cells was used to challenge naive cells expressing the ACE2 receptor. RdRP expression was not detected in naive cells treated with the replicon cell supernatant (the histogram marked as Challenge), while the replicon-verified cells exhibited strong expression of the replicon.

### Establishing the replication capabilities of the replicon

We tested the replication ability of the replicon in several mammalian cells. We transfected HEK293T, BHK21, and VeroE6 cells with three levels of the replicon RNA, namely 2ng, 20ng, and 200ng. As a negative control, we transfected these cells with mRNA of nLuc that can generate a bioluminescence signal similar to our replicon but cannot replicate. In our preliminary experiments, we found widespread cell death within 24 hours post-transfection with the replicon, but not with the nLuc mRNA, providing an initial signal that the replicon induces toxicity, in line with previous SARS-CoV-2 replicon studies^16^. To mitigate this issue, we treated all cells with 100ng/ml of ruxolitinib to inhibit interferon response, which resulted in prolonged cell viability of up to 72hrs.

The expression dynamics of our replicon showed its replication capabilities. We traced the bioluminescence signal of the cells transfected with the replicon for 72 hours and compared them to the bioluminescence of the cells transfected with the FLuc mRNA. Our results show a monotonic increase over time in the ratio between the bioluminescence signal of the replicon compared to the mRNA FLuc in nearly all conditions tested (**Figure 4A; Supp. Table 2**). For instance, after transfecting the HEK293T cells with 2ng of replicon, the initial (24 hours post-transfection) bioluminescence ratio between the replicon and nLuc was only ~0.007x, but it increased to 2x at 72 hours post-transfection. This increase of more than 3 orders of magnitude demonstrates the starkly different dynamics of these two RNA molecules. We found monotonic increases, albeit to a more modest extent, in BHK21 and VeroE6 cells as well. Taken together, our results indicate different decay dynamics between the replicon and a non-replicating mRNA, consistent with a functional replication activity of the replicon.

**Figure 4:**
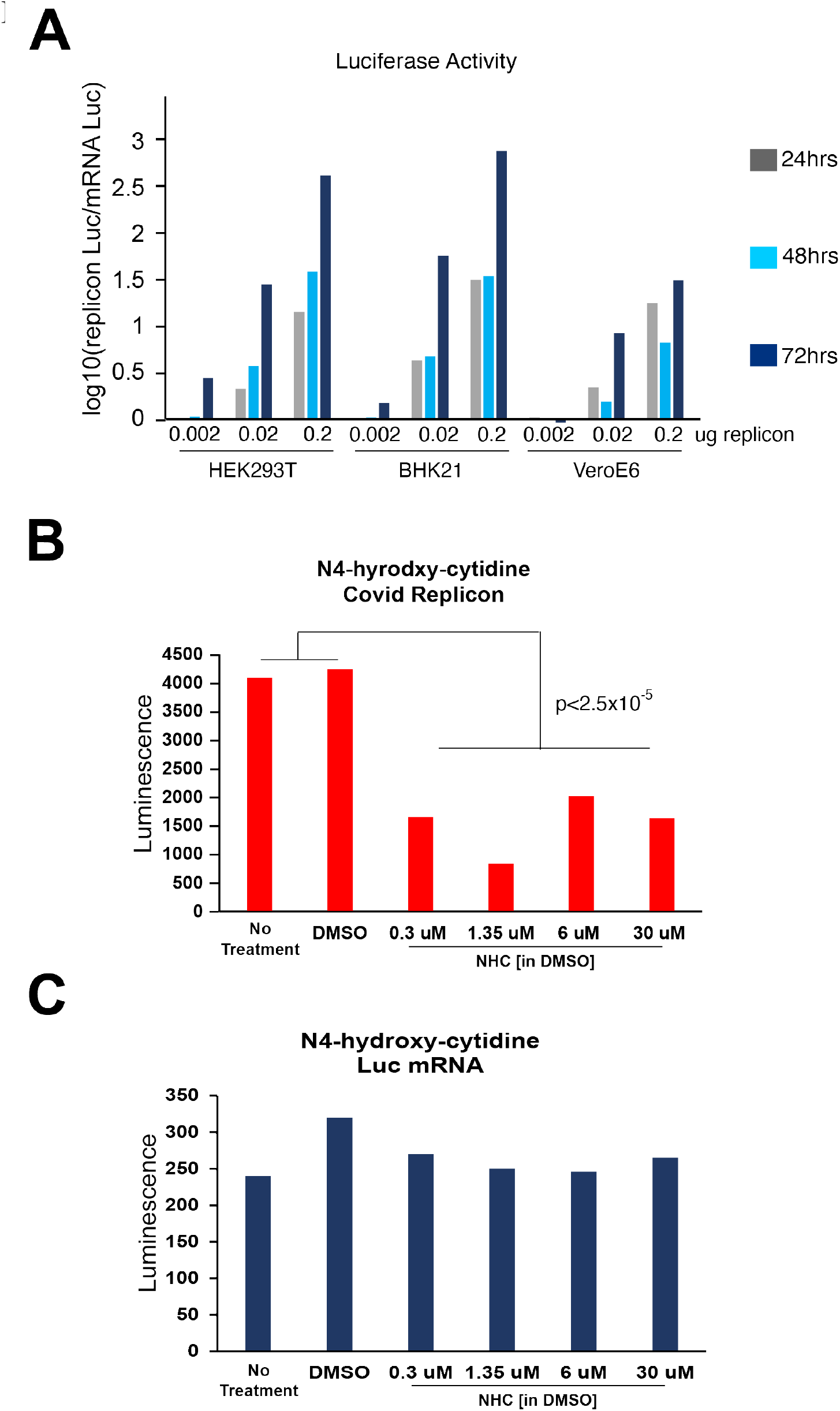
Replication capabilities of the SARS-CoV-2 replicon. **A.** HEK293T, BHK21 and VeroE6 cells were transfected with 0.2ug, 0.02ug and 0.002ug replicon mRNA (capped and polyAdenylated). Luciferase activity was monitored up to 72 hours at 24 hour intervals. **B.** HEK293T cells were transfected with 0.2ug replicon mRNA and the replication activity was challenged with the RdRP inhibitor N4-Hydroxycytidine (Focus Biochemicals). 0.3, 1.35, 6 and 30 uM NHC concentrations were used. *p*-value was calculated with a one-tailed *t-test.* **C.** Luciferase activity both for replicon and Fluc mRNA were measured at 36 hours post transfection.

We further confirmed the replication capabilities of the replicon by subjecting it to N4-Hydroxycytidine (NHC) treatment, the active ingredient in the COVID-19-authorized drug molnupiravir (Merck). NHC is a nucleoside analog that induces G->A mutations, eventually inducing an error catastrophe in several RNA viruses. Importantly, NHC intervenes exclusively in the replication cycle of the virus by the RdRP, and therefore its effect serves as a strong indicator for replication activity. We subjected 250×10^3^ HEK293T replicon-transfected cells to ascending levels of NHC, from 0.3uM to 30uM for 36 hours. In addition, we subjected cells that were transfected with the nLuc mRNA to the same levels of NHC as a negative control. We observed significant (p<2.5×10^-5^) reduction in the bioluminescence signal in the NHC conditions compared to no treatment or mock treatment with DMSO. However, the signal did not monotonically increase in proportion with the concentrations of NHC. The non-replicating mRNA showed no difference in bioluminescence across any of the NHC treatments as compared to no-treatment controls (**Figure 4B-C**). To further support this observation, we examined the GFP signal by microscopy, as well as RdRP gene expression of the SARS-CoV-2 replicon by qPCR. NHC treated HEK293T replicon-transfected cells showed a decline both in GFP signal and in RdRP expression (**Supp. Figure 1**), providing additional support to the self-replicating capabilities of our replicon product and its utility in capturing essential properties of the biology of SARS-CoV-2.

Finally, we demonstrated that our replicon can be specifically suppressed by RNA interference (RNAi). We co-transfected HEK293T cells with 0.2ug of the replicon and one of the following siRNA treatments: (a) S5/S3: a highly efficient siRNA cocktail targeting the subgenomic region of SARS-CoV-2 that was developed by us; (b) J5: an siRNA that was identified in previous studies as a potent inhibitor of HCV, but does not complement the SARS-CoV-2 genome and is therefore a negative control; and (c) no siRNA treatment. To test the specific inhibition of the siRNAs, we sequenced the genome of the replicon system 36 hours posttransfection using the NEBNext VarSkip SARS-CoV-2 amplicon sequencing using Illumina iSeq100 with 150nt PE reads. We then aligned the amplicons to the replicon genome and analyzed the fold change in the number of reads along the sequence across treatment conditions. This analysis showed a 10x-15x decrease in the number of reads about 500nt around the S3 and S5 specific sites, matching our expectation for the RNAi silencing activity against the replicon system (**Figure 5**). Importantly, we did not find any other sharp decrease in coverage of the replicon genome, attesting to the specificity of inhibition. Similarly, we did not observe any decrease in coverage when comparing J5 to no treatment (**Supp. Figure 2**). qPCR analysis of the RdRP region showed a 7% decrease in replicon load when cells were treated with S3/S5 as compared to no treatment. However, we also found a 15% decrease in the replicon load when cells were treated with J5 as compared to no treatment (**Supp. Figure 3**). Since we have not seen any specific reduction in the sequencing coverage of J5, our results suggest a non-specific effect via other mechanisms.

**Figure 5:**
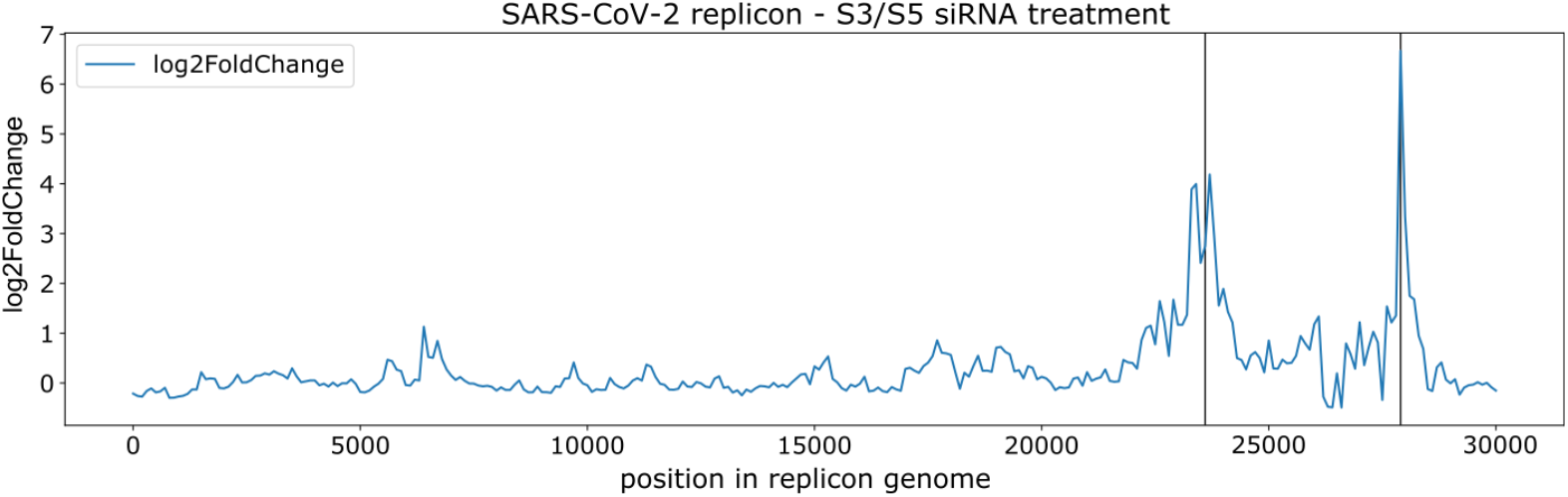
DeSEQ2 analysis of SARS-CoV-2 replicon treatment with S3 and S5 siRNA cocktail. 36 hours post transfection, replicon mRNA from transfected HEK293T cells was sequenced using the NEBNext VarSkip SARS-CoV-2 amplicon kit with Illumina iSeq at 150nt PE reads. We observed a sharp coverage decrease around the S3 (23639-23660 nts) and S5 cleavage (27923-27944 nts) sites (FDR values 4×10^-83^ and 6×10^-22^, respectively) along the replicon sequence.

## Discussion

Replicon systems have been successfully used as mimics of RNA viruses such as Zika, Dengue, and SARS-CoV-1 to facilitate safe and rapid development of novel therapeutics. Here, we reported the development of a safe and efficient SARS-CoV-2 replicon-generating engine, empowered by massively parallel DNA synthesis. The first advantage of our system is that virus infectivity is eliminated while ~97% of its genome is kept intact. Another advantage of our system is its simplicity and ease of use as it only requires the sequence data without access to an actual viral sample. We demonstrated the system with the Beta (B.1.351) variant of SARS-CoV-2 but with its inherent flexibility, it can produce other variants, such as Delta (B.1.617.2) or Omicron (B.1.1.529). We also showed that our replicon precisely captured the genomic sequences of the strain despite utilizing de-novo synthesis and responded to N4-Hydroxycytidine (an RdRP inhibitor) as well as to an siRNA cocktail we developed.

We envision that the ability to rapidly produce widely-available safe replicons is another component in the world’s response to emerging SARS-CoV-2 variants. Using our framework, we can enable researchers to establish a high throughput assay in a BSL-2 lab in three weeks once a new variant is detected. Moreover, by expressing the mutated structural proteins (S, E, and M) in trans, it is possible to produce single-cycle virions to test Spike mutations in the genomic context of SARS-CoV-2^6^. Rapid accessibility of safe viral alternatives can remove critical barriers and unleash more brain power in studying new variants.

Two years of living through the SARS-CoV-2 pandemic taught us that time is of the essence. The field recently developed an automatic monitoring system for emerging SARS-CoV-2 variants^17^. Synthetic biology can complement these efforts by enabling rapid prototyping of therapeutic interventions for variants of concerns.

## Methods

### Bioinformatics pipeline

First, we downloaded all SARS-CoV-2 genomes having Pango lineage B.1.351 from the NCBI virus database (https://www.ncbi.nlm.nih.gov/labs/virus/vssi/#/virus?SeqType_s=Nucleotide&VirusLineage_ss=SARS-CoV-2,%20taxid:2697049&Lineage_s=B.1.351) on June 13 2021 (2,302 sequences). We constructed a multiple sequence alignment using the “-diags” option in MUSCLE v3.8.1551 ^18^. We then computed the consensus sequence using a majority vote. Since the 5’ and 3’ ends of the genome were not preserved in the consensus sequence (because some sequences were missing them), we added the 5’-most 50 and 3’-most 167 nucleotides manually using the reference genome of the SARS-CoV-2 wildtype strain (NCBI reference sequence NC_045512.2). We selected the numbers 50 and 167 based on exact homology between the Wildtype reference genome sequence and the consensus sequence of the 100 nucleotides consecutive to the appended regions. Overall, the consensus sequence included 29,885 nucleotides (vs. 29,903 nucleotides for the SARS-CoV-2 reference genome).

Second, we obtained the reference genome sequence of each SARS-CoV-2 gene (ORF1ab, S, N, E, ORF3a, ORF8, M, ORF7a, ORF6, ORF7b, ORF10) from the reference genome sequence, based on gene coordinates obtained from the NCBI gene database (https://www.ncbi.nlm.nih.gov/gene/?term=NC_045512.2). We used these to find the positions of each gene in the consensus sequence, by finding the 5’ end of each gene in the consensus sequence (based on homology of at least 30 nucleotides with the reference genome sequence) and then translating the resulting protein until hitting a stop codon. We added special handling for the ORF1ab gene, which includes a ribosomal slippage site, by skipping 15 nucleotides at the slippage site (ATCGTTTTTAAACGG) from the translation computation.

Third, we created a modified Spike gene by (1) removing the furin cleavage site (PRRA), and (2) inserting the sequence TAAGTAA (corresponding to a stop codon, a frameshift mutation and a second stop codon) after the first 27 nucleotides.

Fourth, we downloaded the sequence of the synthetic construct clone icSARS-CoV-2-nLuc-GFP (Genbank ID MT461671.1) and extracted the sequence corresponding to the ORF7a-GFP-nLuc fusion protein (nucleotides 27393-28659). We further removed the 5’-most 36 nucleotides, which correspond to ORF7a, from the extracted sequence.

Finally, we constructed the replicon sequence in a 5’->3’ direction by concatenating the following five sequences: (1) All nucleotides in the consensus sequence to the 5’ end of the Spike gene start codon (non-inclusive); (2) the modified Spike gene described above; (3) all nucleotides in the consensus sequence to the 3’ of the Spike gene stop codon and to the 5’ of the E gene start codon (non-inclusive); (4) the modified GFP-nLuc sequence; and (5) all nucleotides in the consensus sequence to the 3’ of the M gene stop-codon.

We performed several diagnostics to verify that the constructed sequence adheres to multiple safety standards and has all the required fail-safe design mechanisms. Specifically, we verified that (1) the 30-nucleotide 5’ and 3’ ends of the E and M genes are missing from the replicon sequence; (2) the Spike protein is only 10 amino acids long; (3) The furin cleavage site is missing from the replicon sequence. As positive controls to our code, we verified that we can find all these genomic elements in the consensus sequence. We additionally verified that we can find the leader transcriptional regulatory sequence (TRS) (ACGAAC) and the body TRS (AAGAAC) in almost the same coordinates in the consensus sequence and the replicon sequence (up to 320 nucleotides apart), except for the TRS corresponding to the removed genes, and with an additional body TRS for the GFP-nLuc gene that we introduced (at position 26,696).

### Replicon Construction

To assemble the 30kb replicon construct, 18 adaptor-on non-clonal gene fragments of roughly 1.6kb were designed with 40bp of homology between fragments and were synthesized by Twist Bioscience. *In vitro* homology based assembly of the 5kb fragments was done by incubating three 1.6kb fragments (40fmol each) with proprietary assembly mix (Twist) at 65°C for 30 minutes in a thermocycler (Invitrogen). The final 5kb product was enriched by PCR with specific primers using Q5 2x MasterMix (NEB). Another round of assembly was repeated by mixing three 5kb fragments to generate 15kb fragments.

The 2 final 15kb fragments were gel extracted via agarose gel electrophoresis and purified with the Large Fragment DNA Recovery Kit (Zymo Research) and eluted in water. Cloning was carried out by adding the two 15kb fragments (80fmol each) to 40fmol pSMART BAC vector (Lucigen) and incubating them with a proprietary assembly mix (Twist) at 65°C for 30 minutes. The reaction was diluted 5x fold and electroporated (Bio-Rad) into Lucigen Replicator V2.0 bacterial cells for outgrowth in LB Lennox Chlor-25 plates (Teknova) overnight at 37°C.

Forty eight colonies were picked for overnight growth, induced with Arabinose for higher copy BAC plasmid, and mini-prepped with QIAprep Spin Miniprep Kit (Qiagen). Plasmids were NGS library prepped with Nextera XT DNA Sample Prep Kit (Illumina) and sequenced 2x×150bp by MiSeq (Illumina). A quarter of the MiSeq run (1.2Gb) was assigned to the 48 samples, yielding roughly 5-6M total reads. NGS analysis was done by alignment with BWA-MEM to the reference and the bam file was visualized on Integrative Genomics Viewer (IGV). Error-free clones were selected post analysis and glycerol banked.

### BAC vector amplification and *in vitro* transcription

To purify BAC plasmids carrying the SARS-CoV-2 replicon, Lucigen Replicator V2.0 bacterial cells were used. Bacterial cultures were induced with Arabinose for multicopy plasmids. Macharey-Nagel NucleoSnap Midi (Fisher Scientific) kit was used for plasmid purification.

Prior to *in vitro* transcription (IVT), BAC plasmids were linearised using AgeI-HF (NEB). Briefly, 4ug plasmid was linearised with 10ul of AgeI at 37°C for 4 hours. The linearised plasmid was precipitated with 2.5x volume of ethanol and 0.1x volume of sodium acetate (pH=5.2) and dissolved in nuclease-free water.

For IVT, the mMachine mMessage kit (Invitrogen) was used with minor modifications. First, the modified GTP analog to naked GTP ratio was set to 1:10 to enable a successful reaction. Additionally, RNase Inhibitor (Promega), 50 mM HEPES pH 7.5, and 10 mM NaCl were used in the final buffer conditions. Reactions were run at 30°C or 27°C for 24 hours. Following DNaseI (Thermofisher, Turbo) treatment, RNA was precipitated with 0.1M LiCl and washed with 70% EtOH three times at 4°C. Precipitated RNA was dissolved in nuclease-free water treated with RNase Inhibitor (Promega). PolyAdenylation was performed using the PolyA Tailing Kit (Thermofisher) per 20ug capped mRNA.

### Transfection

The cells were harvested using TrypLE Express (ThermoFisher), washed with 1X PBS and resuspended in 10ul of electroporation buffer (Neon Transfection Kit, Thermofisher). 0.5 million cells were mixed with 0.2ug, 0.02ug or 0.002ug replicon mRNA (capped and polyAdenylated) diluted in nuclease-free water or with 1ul FLuc mRNA (Trilink). Prog. 7 was used to electroporate the cell-replicon mRNA mixture. The mixture was then immediately transferred to pre-warmed cDMEM (10% FBS), supplemented with 100ng/ml interferon inhibitors (ruxolitinib, SelleckChem) in a 12-well plate. 12 hours later, the media was replaced to remove dead cells arising from electroporation.

### siRNA testing

0.5 million HEK293T cells, 0.2ug replicon and 100nM siRNA duplex were mixed in the electroporation buffer (Neon Transfection Kit, Thermofisher). Prog. 7 was used to electroporate the cell-replicon mixture. The mixture was then immediately transferred to pre-warmed cDMEM (10% FBS), supplemented with 100ng/ml interferon inhibitors (ruxolitinib, SelleckChem) in a 12-well plate. 12 hours later, the media was replaced to remove dead cells arising from electroporation. Cells were harvested 36 hours after transfection.

### Luciferase assay

Cells were harvested in TrypLE Express and resuspended in cDMEM. After centrifugation, the pellet was dissolved in 80ul cDMEM. Luciferase assay was performed using the Dual Luciferase Reporter Assay System kit (Promega). Bioluminescence was measured using CLARIOstar (BMG LABTech).

### Quantitative real time PCR

Quantitative real time PCR (qPCR) was performed using QuantStudio3 (Thermofisher) with Taqman primers as published in the Berlin protocol^15^.

### SARS-CoV-2 amplicon sequencing

RNA was extracted from HEK293T transfected with 0.2ug replicon alone or in combination with 100nM siRNA duplexes (S3/S5 or J5) using Qiagen RNeasy Plus kit.200ng of extracted RNA was used to produce SARS-CoV-2 amplicon libraries using the NEBNext SARS-CoV-2 FS Library Prep Kit (Illumina) using the VarSkip Short Express Protocol with 25 minutes fragmentation step and 8 cycles of PCR enrichment. The libraries were multiplexed using the NEBNext Multiplex Oligos for Illumina (Index Primers Set 1) and sequenced paired end 150 using an Illumina iSEQ100.

### Replicon coverage pipeline

We trimmed the adapters of pair-end reads with minimal length of 5 bases using cutadapt v3.5^19^ and mapped the remaining reads to the replicon reference using Bowtie2 v2.4.4^20^. Next, we converted the SAM output files to sorted BAM files using SAMtools v1.11^21^ and extracted the coverage as Wig files using the IGVTools^22^ count function, with a window size of 1 for reads with a minimal mapping quality of 1.

In order to conduct differential expression analysis, we partitioned the replicon genome into non-overlapping tiles of 100 bases. For each sample, we set each tile expression to be the maximal coverage within that range. We used DESeq2^23^ for statistical analysis. The replicon and replicon plus J5 samples served as controls against the replicon plus S3/S5 sample. Since the siRNA degraded a large proportion of the replicon genome, we calibrated the null distribution of DESeq2 using the control genes parameter, using the first 200 tiles (corresponding to the first 20,000 bases of the replicon genome). The first 200 windows are relatively distant from the siRNA target sites, and therefore should not be affected by possible siRNA cleavage

## Supplementary Figures

### Supplementary Figure 1

**Supp. Figure 1:**
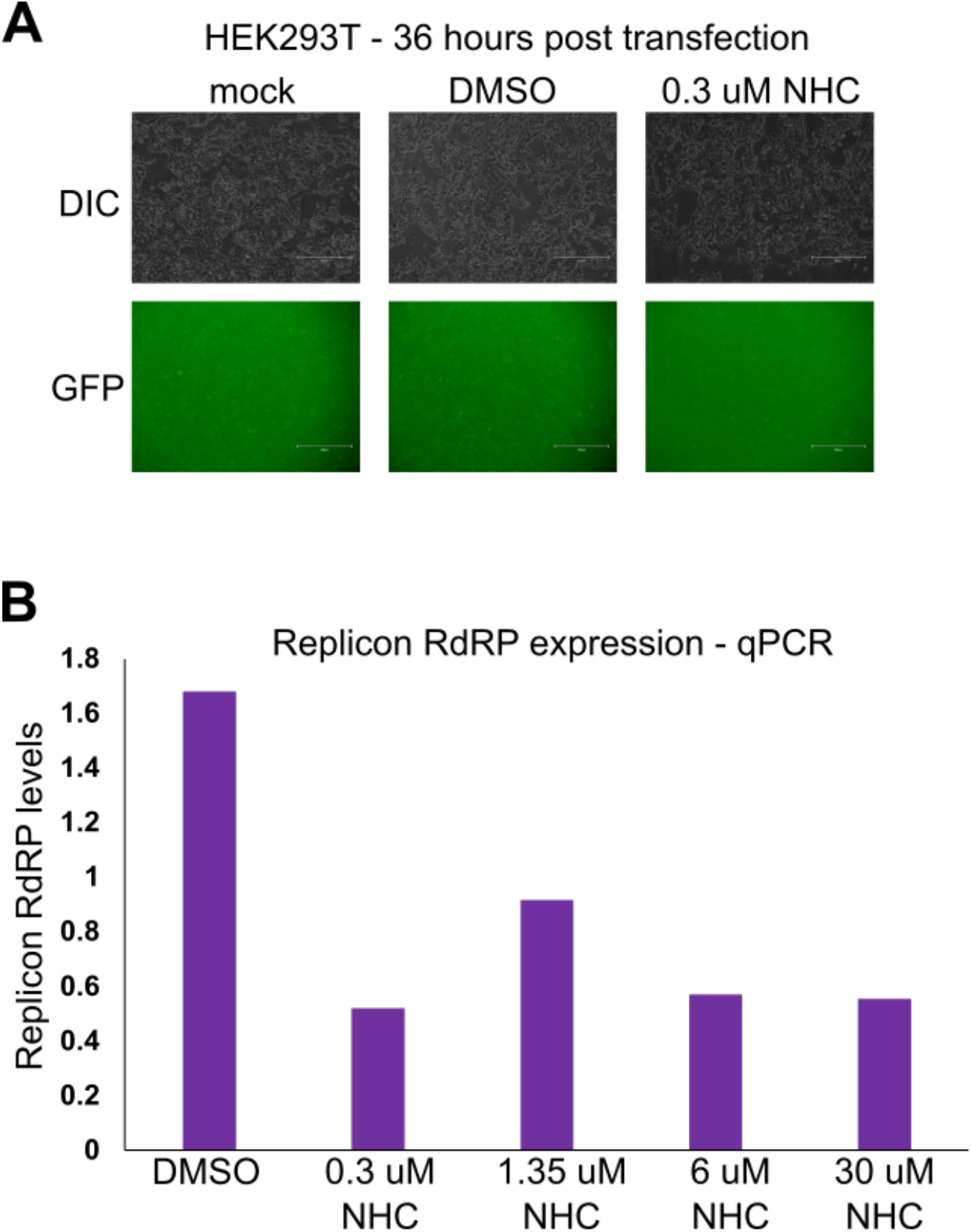
Testing the impact of a clinically authorized drug on the replicon. **A.** 0.3 uM N4-Hydroxycytidine (NHC, for which molnupiravir is a prodrug) inhibits the GFP signal of the SARS-CoV-2 replicon. GFP images were taken 36 hours post transfection using Evos microscope (Invitrogen) under 10X magnification. **B.** RdRP expression of the replicon upon treatment with 0.3uM, 1.35uM, 6uM and 30 uM NHC concentrations was performed using QuantStudio3. RdRP expression declines upon treatment with all concentrations of NHC, normalized to no treatment conditions.

### Supplementary Figure 2

**Supp. Figure 2:**
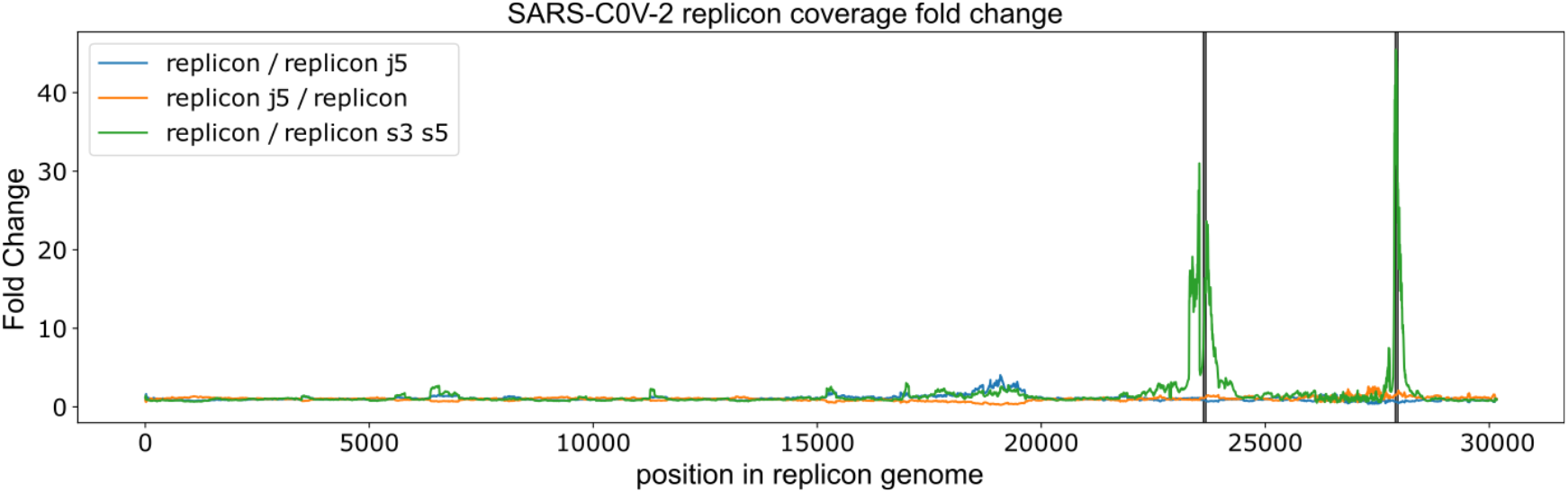
Replicon coverage fold change compared to S3/S5 and J5 siRNAs using raw coverage reads. We observed a sharp decrease in the replicon when treated with S3/S5 siRNA cocktail. However, as expected, no decrease was observed for the negative control (J5 siRNA treatment). Unlike Figure 5, this figure is based on raw coverage readouts.

### Supplementary Figure 3

**Supp. Figure 3:**
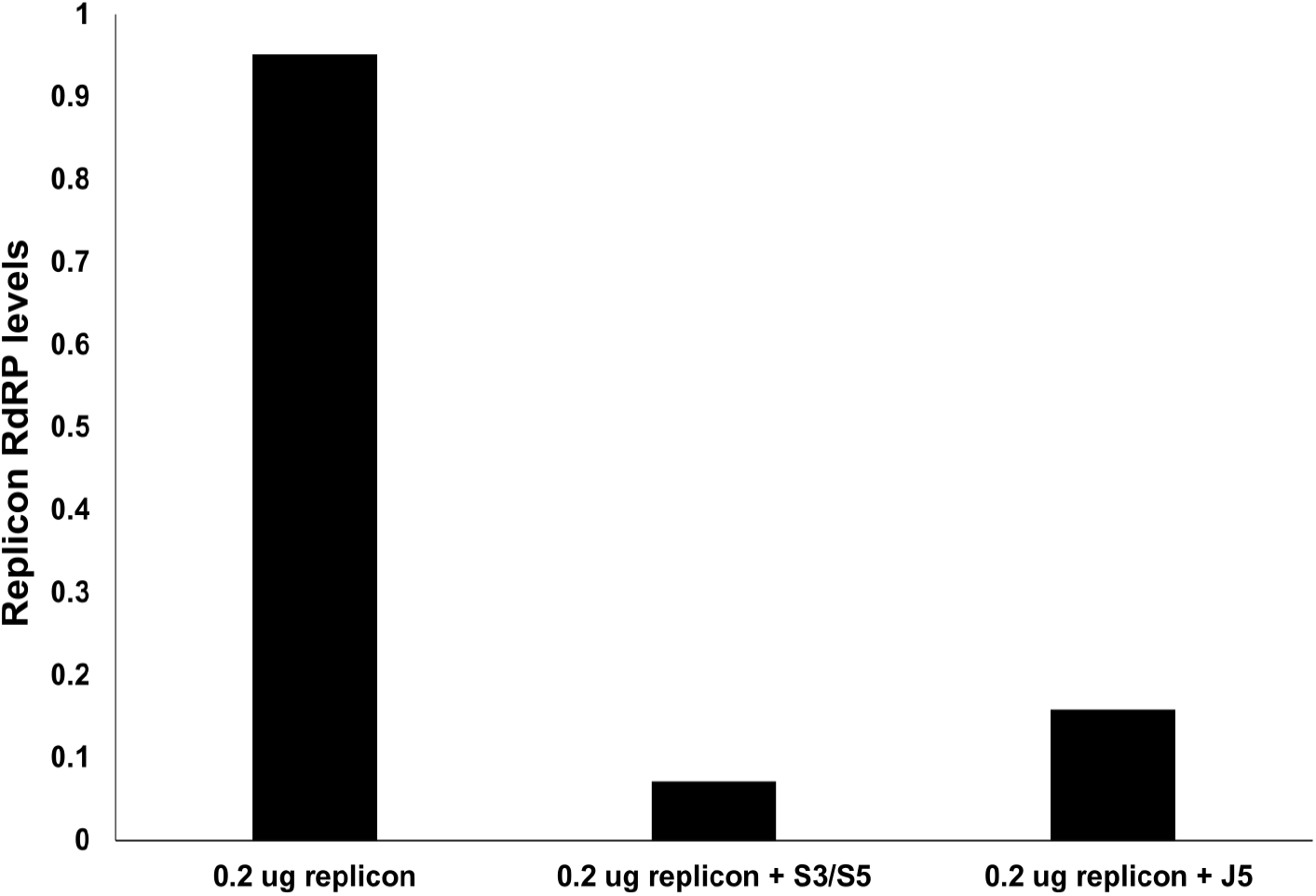
qPCR levels of the replicon RdRP following siRNA treatments, normalized to no treatment conditions.

## Supplementary Tables

### Supplementary Table 1

**Supp. Table 1:**
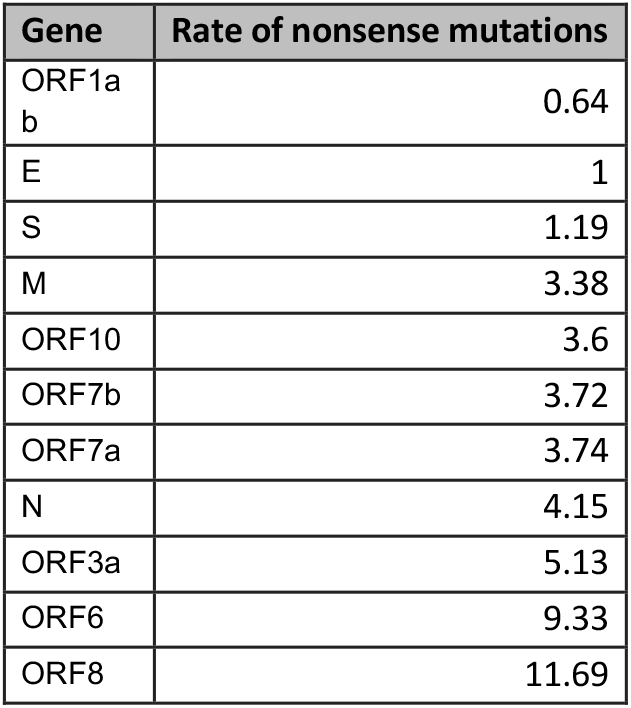
The rate of nonsense mutation per nucleotide in 26,000 genomes of SARS-CoV2 normalized to the rate of nonsense mutations in the E gene.

| Gene | Rate of nonsense mutations |
| --- | --- |
| ORF1a<br>b | 0.64 |
| E | 1 |
| S | 1.19 |
| M | 3.38 |
| ORF10 | 3.6 |
| ORF7b | 3.72 |
| ORF7a | 3.74 |
| N | 4.15 |
| ORF3a | 5.13 |
| ORF6 | 9.33 |
| ORF8 | 11.69 |

### Supplementary Table 2

**Supp Table 2:** The raw values of bioluminescence from each condition compared to non-replicating luciferase (luciferase control).

| cells | treatment | time point |  |  |  |  |  |
| --- | --- | --- | --- | --- | --- | --- | --- |
| HEK293T |  | 24 hrs |  | 48 hrs |  | 72 hrs |  |
|  |  | rep1 | rep2 | rep1 | rep2 | rep1 | rep2 |
|  | 0.2ug | 5070 | 4380 | 5400 | 3880 | 1200 | 6600 |
|  | 0.02ug | 470 | 320 | 440 | 220 | 220 | 280 |
|  | 0.002ug | 0 | 5 | 14 | 2 | 16 | 16 |
|  | GFP control | 7 | 7 | 6 | 8 | 6 | 20 |
|  | Luciferase control | 200 | 500 | 120 | 120 | 0 | 18 |
|  | Mock | 7 | 0 | 6 | 2 | 2 | 8 |
| BHK21 |  | 24 hrs |  | 48 hrs |  | 72 hrs |  |
|  |  | rep1 | rep2 | rep1 | rep2 | rep1 | rep2 |
|  | 0.2ug | 5770 | 6640 | 6656 | 2888 | 3590 | 2800 |
|  | 0.02ug | 400 | 910 | 340 | 720 | 180 | 280 |
|  | 0.002ug | 7 | 4 | 8 | 4 | -2 | 6 |
|  | GFP control | 3 | 4 | 2 | 0 | 2 | 16 |
|  | Luciferase control | 200 | 200 | 160 | 120 | 0 | 8 |
|  | Mock | 1 | 1 | 0 | 0 | 4 | -2 |
| VeroE6 |  | 24 hrs |  | 48 hrs |  | 72 hrs |  |
|  |  | rep1 | rep2 | rep1 | rep2 | rep1 | rep2 |
|  | 0.2ug | 1710 | 1680 | 560 | 1160 | 600 | 140 |
|  | 0.02ug | 190 | 50 | 40 | 120 | 40 | 140 |
|  | 0.002ug | 3 | 5 | 0 | 0 | -2 | 0 |
|  | GFP control | 8 | 1 | 4 | 2 | 4 | 10 |
|  | Luciferase control | 0 | 200 | 180 | 120 | 4 | 20 |
|  | Mock | 4 | 5 | 6 | 6 | 8 | 12 |

## Supplemental Data File 1

**>eleven_bl_3 5 l_gfp_luc_TWIST**

ATTAAAGGTTTATACCTTCCCAGGTAACAAACCAACCAACTTTCGATCTCTTGTAGATCTGTTCTCTAAACGAACTTTAAAATCTGTGTGGCTGTCACTCGGCTGCATGCTTAGTGCACTCACGCAGTATAATTAATAACTAATTACTGTCGTTGACAGGACACGAGTAACTCTTCTATCTTCTGCAGGCTGCTTACGGTCTCGTCCGTGTTGCAGCCGATCATCAGCACATCTAGGTTTTGTCCGGGTGTGACCGAAAGGTAAGATGGAGAGCCTTGTCCCTGGTTTCAACGAGAAAACACACGTCCAACTCAGTTTGCCTGTTTTACAGGTTCGCGACGTGCTCGTACGTGGCTTTGGAGACTCCGTGGAGGAGGTCTTATCAGAGGCACGTCAACATCTTAAAGATGGCACTTGTGGCTTAGTAGAAGTTGAAAAAGGCGTTTTGCCTCAACTTGAACAGCCCTATGTGTTCATCAAACGTTCGGATGCTCGAACTGCACCTCATGGTCATGTTATGGTTGAGCTGGTAGCAGAACTCGAAGGCATTCAGTACGGTCGTAGTGGTGAGACACTTGGTGTCCTTGTCCCTCATGTGGGCGAAATACCAGTGGCTTACCGCAAGGTTCTTCTTCGTAAGAACGGTAATAAAGGAGCTGGTGGCCATAGTTACGGCGCCGATCTAAAGTCATTTGACTTAGGCGACGAGCTTGGCACTGATCCTTATGAAGATTTTCAAGAAAACTGGAACACTAAACATAGCAGTGGTGTTACCCGTGAACTCATGCGTGAGCTTAACGGAGGGGCATACACTCGCTATGTCGATAACAACTTCTGTGGCCCTGATGGCTACCCTCTTGAGTGCATTAAAGACCTTCTAGCACGTGCTGGTAAAGCTTCATGCACTTTGTCCGAACAACTGGACTTTATTGACACTAAGAGGGGTGTATACTGCTGCCGTGAACATGAGCATGAAATTGCTTGGTACACGGAACGTTCTGAAAAGAGCTATGAATTGCAGACACCTTTTGAAATTAAATTGGCAAAGAAATTTGACATCTTCAATGGGGAATGTCCAAATTTTGTATTTCCCTTAAATTCCATAATCAAGACTATTCAACCAAGGGTTGAAAAGAAAAAGCTTGATGGCTTTATGGGTAGAATTCGATCTGTCTATCCAGTTGCGTCACCAAATGAATGCAACCAAATGTGCCTTTCAACTCTCATGAAGTGTGATCATTGTGGTGAAACTTCATGGCAGACGGGCGATTTTGTTAAAGCCACTTGCGAATTTTGTGGCACTGAGAATTTGACTAAAGAAGGTGCCACTACTTGTGGTTACTTACCCCAAAATGCTGTTGTTAAAATTTATTGTCCAGCATGTCACAATTCAGAAGTAGGACCTGAGCATAGTCTTGCCGAATACCATAATGAATCTGGCTTGAAAACCATTCTTCGTAAGGGTGGTCGCACTATTGCCTTTGGAGGCTGTGTGTTCTCTTATGTTGGTTGCCATAACAAGTGTGCCTATTGGGTTCCACGTGCTAGCGCTAACATAGGTTGTAACCATACAGGTGTTGTTGGAGAAGGTTCCGAAGGTCTTAATGACAACCTTCTTGAAATACTCCAAAAAGAGAAAGTCAACATCAATATTGTTGGTGACTTTAAACTTAATGAAGAGATCGCCATTATTTTGGCATCTTTTTCTGCTTCCACAAGTGCTTTTGTGGAAACTGTGAAAGGTTTGGATTATAAAGCATTCAAACAAATTGTTGAATCCTGTGGTAATTTTAAAGTTACAAAAGGAAAAGCTAAAAAAGGTGCCTGGAATATTGGTGAACAGAAATCAATACTGAGTCCTCTTTATGCATTTGCATCAGAGGCTGCTCGTGTTGTACGATCAATTTTCTCCCGCACTCTTGAAACTGCTCAAAATTCTGTGCGTGTTTTACAGAAGGCCGCTATAACAATACTAGATGGAATTTCACAGTATTCACTGAGACTCATTGATGCTATGATGTTCACATCTGATTTGGCTACTAACAATCTAGTTGTAATGGCCTACATTACAGGTGGTGTTGTTCAGTTGACTTCGCAGTGGCTAACTAACATCTTTGGCACTGTTTATGAAAAACTCAAACCCGTCCTTGATTGGCTTGAAGAGAAGTTTAAGGAAGGTGTAGAGTTTCTTAGAGACGGTTGGGAAATTGTTAAATTTATCTCAACCTGTGCTTGTGAAATTGTCGGTGGACAAATTGTCACCTGTGCAAAGGAAATTAAGGAGAGTGTTCAGACATTCTTTAAGCTTGTAAATAAATTTTTGGCTTTGTGTGCTGACTCTATCATTATTGGTGGAGCTAAACTTAAAGCCTTGAATTTAGGTGAAACATTTGTCACGCACTCAAAGGGATTGTACAGAAAGTGTGTTAAATCCAGAGAAGAAACTGGCCTACTCATGCCTCTAAAAGCCCCAAAAGAAATTATCTTCTTAGAGGGAGAAACACTTCCCACAGAAGTGTTAACAGAGGAAGTTGTCTTGAAAACTGGTGATTTACAACCATTAGAACAACCTACTAGTGAAGCTGTTGAAGCTCCATTGGTTGGTACACCAGTTTGTATTAACGGGCTTATGTTGCTCGAAATCAAAGACACAGAAAAGTACTGTGCCCTTGCACCTAATATGATGGTAACTAACAATACCTTCACACTCAAAGGCGGTGCACCAACAAAGGTTACTTTTGGTGATGACACTGTGATAGAAGTGCAAGGTTACAAGAGTGTGAATATCACTTTTGAACTTGATGAAAGGATTGATAAAGTACTTAATGAGAAGTGCTCTGCCTATACAGTTGAACTCGGTACAGAAGTAAATGAGTTCGCCTGTGTTGTGGCAGATGCTGTCATAAAAACTTTGCAACCAGTATCTGAATTACTTACACCACTGGGCATTGATTTAGATGAGTGGAGTATGGCTACATACTACTTATTTGATGAGTCTGGTGAGTTTAAATTGGCTTCACATATGTATTGTTCTTTTTACCCTCCAGATGAGGATGAAGAAGAAGGTGATTGTGAAGAAGAAGAGTTTGAGCCATCAACTCAATATGAGTATGGTACTGAAGATGATTACCAAGGTAAACCTTTGGAATTTGGTGCCACTTCTGCTGCTCTTCAACCTGAAGAAGAGCAAGAAGAAGATTGGTTAGATGATGATAGTCAACAAACTGTTGGTCAACAAGACGGCAGTGAGGACAATCAGACAACTACTATTCAAACAATTGTTGAGGTTCAACCTCAATTAGAGATGGAACTTACACCAGTTGTTCAGACTATTGAAGTGAATAGTTTTAGTGGTTATTTAAAACTTACTGACAATGTATACATTAAAAATGCAGACATTGTGGAAGAAGCTAAAAAGGTAAAACCAACAGTGGTTGTTAATGCAGCCAATGTTTACCTTAAACATGGAGGAGGTGTTGCAGGAGCCTTAAATAAGGCTACTAACAATGCCATGCAAGTTGAATCTGATGATTACATAGCTACTAATGGACCACTTAAAGTGGGTGGTAGTTGTGTTTTAAGCGGACACAATCTTGCTAAACACTGTCTTCATGTTGTCGGCCCAAATGTTAACAAAGGTGAAGACATTCAACTTCTTAAGAGTGCTTATGAAAATTTTAATCAGCACGAAGTTCTACTTGCACCATTATTATCAGCTGGTATTTTTGGTGCTGACCCTATACATTCTTTAAGAGTTTGTGTAGATACTGTTCGCACAAATGTCTACTTAGCTGTCTTTGATAAAAATCTCTATGACAAACTTGTTTCAAGCTTTTTGGAAATGAAGAGTGAAAAGCAAGTTGAACAAAAGATCGCTGAGATTCCTAAAGAGGAAGTTAAGCCATTTATAACTGAAAGTAAACCTTCAGTTGAACAGAGAAAACAAGATGATAAGAAAATCAAAGCTTGTGTTGAAGAAGTTACAACAACTCTGGAAGAAACTAAGTTCCTCACAGAAAACTTGTTACTTTATATTGACATTAATGGCAATCTTCATCCAGATTCTGCCACTCTTGTTAGTGACATTGACATCACTTTCTTAAAGAAAGATGCTCCATATATAGTGGGTGATGTTGTTCAAGAGGGTGTTTTAACTGCTGTGGTTATACCTACTAAAAAGGCTGGTGGCACTACTGAAATGCTAGCGAAAGCTTTGAGAAAAGTGCCAACAGACAATTATATAACCACTTACCCGGGTCAGGGTTTAAATGGTTACACTGTAGAGGAGGCAAAGACAGTGCTTAAAAAGTGTAAAAGTGCCTTTTACATTCTACCATCTATTATCTCTAATGAGAAGCAAGAAATTCTTGGAACTGTTTCTTGGAATTTGCGAGAAATGCTTGCACATGCAGAAGAAACACGCAAATTAATGCCTGTCTGTGTGGAAACTAAAGCCATAGTTTCAACTATACAGCGTAAATATAAGGGTATTAAAATACAAGAGGGTGTGGTTGATTATGGTGCTAGATTTTACTTTTACACCAGTAAAACAACTGTAGCGTCACTTATCAACACACTTAACGATCTAAATGAAACTCTTGTTACAATGCCACTTGGCTATGTAACACATGGCTTAAATTTGGAAGAAGCTGCTCGGTATATGAGATCTCTCAAAGTGCCAGCTACAGTTTCTGTTTCTTCACCTGATGCTGTTACAGCGTATAATGGTTATCTTACTTCTTCTTCTAAAACACCTGAAGAACATTTTATTGAAACCATCTCACTTGCTGGTTCCTATAAAGATTGGTCCTATTCTGGACAATCTACACAACTAGGTATAGAATTTCTTAAGAGAGGTGATAAAAGTGTATATTACACTAGTAATCCTACCACATTCCACCTAGATGGTGAAGTTATCACCTTTGACAATCTTAAGACACTTCTTTCTTTGAGAGAAGTGAGGACTATTAAGGTGTTTACAACAGTAGACAACATTAACCTCCACACGCAAGTTGTGGACATGTCAATGACATATGGACAACAGTTTGGTCCAACTTATTTGGATGGAGCTGATGTTACTAAAATAAAACCTCATAATTCACATGAAGGTAAAACATTTTATGTTTTACCTAATGATGACACTCTACGTGTTGAGGCTTTTGAGTACTACCACACAACTGATCCTAGTTTTCTGGGTAGGTACATGTCAGCATTAAATCACACTAAAAATTGGAAATACCCACAAGTTAATGGTTTAACTTCTATTAAATGGGCAGATAACAACTGTTATCTTGCCACTGCATTGTTAACACTCCAACAAATAGAGTTGAAGTTTAATCCACCTGCTCTACAAGATGCTTATTACAGAGCAAGGGCTGGTGAAGCTGCTAACTTTTGTGCACTTATCTTAGCCTACTGTAATAAGACAGTAGGTGAGTTAGGTGATGTTAGAGAAACAATGAGTTACTTGTTTCAACATGCCAATTTAGATTCTTGCAAAAGAGTCTTGAACGTGGTGTGTAAAACTTGTGGACAACAGCAGACAACCCTTAAGGGTGTAGAAGCTGTTATGTACATGGGCACACTTTCTTATGAACAATTTAAGAAAGGTGTTCAGATACCTTGTACGTGTGGTAAACAAGCTACAAAATATCTAGTACAACAGGAGTCACCTTTTGTTATGATGTCAGCACCACCTGCTCAGTATGAACTTAAGCATGGTACATTTACTTGTGCTAGTGAGTACACTGGTAATTACCAGTGTGGTCACTATAAACATATAACTTCTAAAGAAACTTTGTATTGCATAGACGGTGCTTTACTTACAAAGTCCTCAGAATACAAAGGTCCTATTACGGATGTTTTCTACAAAGAAAACAGTTACACAACAACCATAAAACCAGTTACTTATAAATTGGATGGTGTTGTTTGTACAGAAATTGACCCTAAGTTGGACAATTATTATAAGAAAGACAATTCTTATTTCACAGAGCAACCAATTGATCTTGTACCAAACCAACCATATCCAAACGCAAGCTTCGATAATTTTAAGTTTGTATGTGATAATATCAAATTTGCTGATGATTTAAACCAGTTAACTGGTTATAAGAAACCTGCTTCAAGAGAGCTTAAAGTTACATTTTTCCCTGACTTAAATGGTGATGTGGTGGCTATTGATTATAAACACTACACACCCTCTTTTAAGAAAGGAGCTAAATTGTTACATAAACCTATTGTTTGGCATGTTAACAATGCAACTAATAAAGCCACGTATAAACCAAATACCTGGTGTATACGTTGTCTTTGGAGCACAAAACCAGTTGAAACATCAAATTCGTTTGATGTACTGAAGTCAGAGGACGCGCAGGGAATGGATAATCTTGCCTGCGAAGATCTAAAACCAGTCTCTGAAGAAGTAGTGGAAAATCCTACCATACAGAAAGACGTTCTTGAGTGTAATGTGAAAACTACCGAAGTTGTAGGAGACATTATACTTAAACCAGCAAATAATAGTTTAAAAATTACAGAAGAGGTTGGCCACACAGATCTAATGGCTGCTTATGTAGACAATTCTAGTCTTACTATTAAGAAACCTAATGAATTATCTAGAGTATTAGGTTTGAAAACCCTTGCTACTCATGGTTTAGCTGCTGTTAATAGTGTCCCTTGGGATACTATAGCTAATTATGCTAAGCCTTTTCTTAACAAAGTTGTTAGTACAACTACTAACATAGTTACACGGTGTTTAAACCGTGTTTGTACTAATTATATGCCTTATTTCTTTACTTTATTGCTACAATTGTGTACTTTTACTAGAAGTACAAATTCTAGAATTAAAGCATCTATGCCGACTACTATAGCAAAGAATACTGTTAAGAGTGTCGGTAAATTTTGTCTAGAGGCTTCATTTAATTATTTGAAGTCACCTAATTTTTCTAAACTGATAAATATTATAATTTGGTTTTTACTATTAAGTGTTTGCCTAGGTTCTTTAATCTACTCAACCGCTGCTTTAGGTGTTTTAATGTCTAATTTAGGCATGCCTTCTTACTGTACTGGTTACAGAGAAGGCTATTTGAACTCTACTAATGTCACTATTGCAACCTACTGTACTGGTTCTATACCTTGTAGTGTTTGTCTTAGTGGTTTAGATTCTTTAGACACCTATCCTTCTTTAGAAACTATACAAATTACCATTTCATCTTTTAAATGGGATTTAACTGCTTTTGGCTTAGTTGCAGAGTGGTTTTTGGCATATATTCTTTTCACTAGGTTTTTCTATGTACTTGGATTGGCTGCAATCATGCAATTGTTTTTCAGCTATTTTGCAGTACATTTTATTAGTAATTCTTGGCTTATGTGGTTAATAATTAATCTTGTACAAATGGCCCCGATTTCAGCTATGGTTAGAATGTACATCTTCTTTGCATCATTTTATTATGTATGGAAAAGTTATGTGCATGTTGTAGACGGTTGTAATTCATCAACTTGTATGATGTGTTACAAACGTAATAGAGCAACAAGAGTCGAATGTACAACTATTGTTAATGGTGTTAGAAGGTCCTTTTATGTCTATGCTAATGGAGGTAAAGGCTTTTGCAAACTACACAATTGGAATTGTGTTAATTGTGATACATTCTGTGCTGGTAGTACATTTATTAGTGATGAAGTTGCGAGAGACTTGTCACTACAGTTTAAAAGACCAATAAATCCTACTGACCAGTCTTCTTACATCGTTGATAGTGTTAC AGTGAAGAATGGTTCCATCCATCTTTACTTTGATAAAGCTGGTCAAAAGACTTATGAAAGACATTCTCTCTCTCATTTTGTTAACTTAGACAACCTGAGAGCTAATAACACTAAAGGTTCATTGCCTATTAATGTTATAGTTTTTGATGGTAAATCAAAATGTGAAGAATCATCTGCAAAATCAGCGTCTGTTTACTACAGTCAGCTTATGTGTCAACCTATACTGTTACTAGATCAGGCATTAGTGTCTGATGTTGGTGATAGTGCGGAAGTTGCAGTTAAAATGTTTGATGCTTACGTTAATACGTTTTCATCAACTTTTAACGTACCAATGGAAAAACTCAAAACACTAGTTGCAACTGCAGAAGCTGAACTTGCAAAGAATGTGTCCTTAGACAATGTCTTATCTACTTTTATTTCAGCAGCTCGGCAAGGGTTTGTTGATTCAGATGTAGAAACTAAAGATGTTGTTGAATGTCTTAAATTGTCACATCAATCTGACATAGAAGTTACTGGCGATAGTTGTAATAACTATATGCTCACCTATAACAAAGTTGAAAACATGACACCCCGTGACCTTGGTGCTTGTATTGACTGTAGTGCGCGTCATATTAATGCGCAGGTAGCAAAAAGTCACAACATTGCTTTGATATGGAACGTTAAAGATTTCATGTCATTGTCTGAACAACTACGAAAACAAATACGTAGTGCTGCTAAAAAGAATAACTTACCTTTTAAGTTGACATGTGCAACTACTAGACAAGTTGTTAATGTTGTAACAACAAAGATAGCACTTAAGGGTGGTAAAATTGTTAATAATTGGTTGAAGCAGTTAATTAAAGTTACACTTGTGTTCCTTTTTGTTGCTGCTATTTTCTATTTAATAACACCTGTTCATGTCATGTCTAAACATACTGACTTTTCAAGTGAAATCATAGGATACAAGGCTATTGATGGTGGTGTCACTCGTGACATAGCATCTACAGATACTTGTTTTGCTAACAAACATGCTGATTTTGACACATGGTTTAGCCAGCGTGGTGGTAGTTATACTAATGACAAAGCTTGCCCATTGATTGCTGCAGTCATAACAAGAGAAGTGGGTTTTGTCGTGCCTGGTTTGCCTGGCACGATATTACGCACAACTAATGGTGACTTTTTGCATTTCTTACCTAGAGTTTTTAGTGCAGTTGGTAACATCTGTTACACACCATCAAAACTTATAGAGTACACTGACTTTGCAACATCAGCTTGTGTTTTGGCTGCTGAATGTACAATTTTTAAAGATGCTTCTGGTAAGCCAGTACCATATTGTTATGATACCAATGTACTAGAAGGTTCTGTTGCTTATGAAAGTTTACGCCCTGACACACGTTATGTGCTCATGGATGGCTCTATTATTCAATTTCCTAACACCTACCTTGAAGGTTCTGTTAGAGTGGTAACAACTTTTGATTCTGAGTACTGTAGGCACGGCACTTGTGAAAGATCAGAAGCTGGTGTTTGTGTATCTACTAGTGGTAGATGGGTACTTAACAATGATTATTACAGATCTTTACCAGGAGTTTTCTGTGGTGTAGATGCTGTAAATTTACTTACTAATATGTTTACACCACTAATTCAACCTATTGGTGCTTTGGACATATCAGCATCTATAGTAGCTGGTGGTATTGTAGCTATCGTAGTAACATGCCTTGCCTACTATTTTATGAGGTTTAGAAGAGCTTTTGGTGAATACAGTCATGTAGTTGCCTTTAATACTTTACTATTCCTTATGTCATTCACTGTACTCTGTTTAACACCAGTTTACTCATTCTTACCTGGTGTTTATTCTGTTATTTACTTGTACTTGACATTTTATCTTACTAATGATGTTTCTTTTTTAGCACATATTCAGTGGATGGTTATGTTCACACCTTTAGTACCTTTCTGGATAACAATTGCTTATATCATTTGTATTTCCACAAAGCATTTCTATTGGTTCTTTAGTAATTACCTAAAGAGACGTGTAGTCTTTAATGGTGTTTCCTTTAGTACTTTTGAAGAAGCTGCGCTGTGCACCTTTTTGTTAAATAAAGAAATGTATCTAAAGTTGCGTAGTGATGTGCTATTACCTCTTACGCAATATAATAGATACTTAGCTCTTTATAATAAGTACAAGTATTTTAGTGGAGCAATGGATACAACTAGCTACAGAGAAGCTGCTTGTTGTCATCTCGCAAAGGCTCTCAATGACTTCAGTAACTCAGGTTCTGATGTTCTTTACCAACCACCACAAACCTCTATCACCTCAGCTGTTTTGCAGAGTGGTTTTAGAAAAATGGCATTCCCATCTGGTAAAGTTGAGGGTTGTATGGTACAAGTAACTTGTGGTACAACTACACTTAACGGTCTTTGGCTTGATGACGTAGTTTACTGTCCAAGACATGTGATCTGCACCTCTGAAGACATGCTTAACCCTAATTATGAAGATTTACTCATTCGTAAGTCTAATCATAATTTCTTGGTACAGGCTGGTAATGTTCAACTCAGGGTTATTGGACATTCTATGCAAAATTGTGTACTTAAGCTTAGGGTTGATACAGCCAATCCTAAGACACCTAAGTATAAGTTTGTTCGCATTCAACCAGGACAGACTTTTTCAGTGTTAGCTTGTTACAATGGTTCACCATCTGGTGTTTACCAATGTGCTATGAGGCCCAATTTCACTATTAAGGGTTCATTCCTTAATGGTTCATGTGGTAGTGTTGGTTTTAACATAGATTATGACTGTGTCTCTTTTTGTTACATGCACCATATGGAATTACCAACTGGAGTTCATGCTGGCACAGACTTAGAAGGTAACTTTTATGGACCTTTTGTTGACAGGCAAACAGCACAAGCAGCTGGTACGGACACAACTATTACAGTTAATGTTTTAGCTTGGTTGTACGCTGCTGTTATAAATGGAGACAGGTGGTTTCTCAATCGATTTACCACAACTCTTAATGACTTTAACCTTGTGGCTATGAAGTACAATTATGAACCTCTAACACAAGACCATGTTGACATACTAGGACCTCTTTCTGCTCAAACTGGAATTGCCGTTTTAGATATGTGTGCTTCATTAAAAGAATTACTGCAAAATGGTATGAATGGACGTACCATATTGGGTAGTGCTTTATTAGAAGATGAATTTACACCTTTTGATGTTGTTAGACAATGCTCAGGTGTTACTTTCCAAAGTGCAGTGAAAAGAACAATCAAGGGTACACACCACTGGTTGTTACTCACAATTTTGACTTCACTTTTAGTTTTAGTCCAGAGTACTCAATGGTCTTTGTTCTTTTTTTTGTATGAAAATGCCTTTTTACCTTTTGCTATGGGTATTATTGCTATGTCTGCTTTTGCAATGATGTTTGTCAAACATAAGCATGCATTTCTCTGTTTGTTTTTGTTACCTTCTCTTGCCACTGTAGCTTATTTTAATATGGTCTATATGCCTGCTAGTTGGGTGATGCGTATTATGACATGGTTGGATATGGTTGATACTAGTTTGAAGCTAAAAGACTGTGTTATGTATGCATCAGCTGTAGTGTTACTAATCCTTATGACAGCAAGAACTGTGTATGATGATGGTGCTAGGAGAGTGTGGACACTTATGAATGTCTTGACACTCGTTTATAAAGTTTATTATGGTAATGCTTTAGATCAAGCCATTTCCATGTGGGCTCTTATAATCTCTGTTACTTCTAACTACTCAGGTGTAGTTACAACTGTCATGTTTTTGGCCAGAGGTATTGTTTTTATGTGTGTTGAGTATTGCCCTATTTTCTTCATAACTGGTAATACACTTCAGTGTATAATGCTAGTTTATTGTTTCTTAGGCTATTTTTGTACTTGTTACTTTGGCCTCTTTTGTTTACTCAACCGCTACTTTAGACTGACTCTTGGTGTTTATGATTACTTAGTTTCTACACAGGAGTTTAGATATATGAATTCACAGGGACTACTCCCACCCAAGAATAGCATAGATGCCTTCAAACTCAACATTAAATTGTTGGGTGTTGGTGGCAAACCTTGTATCAAAGTAGCCACTGTACAGTCTAAAATGTCAGATGTAAAGTGCACATCAGTAGTCTTACTCTCAGTTTTGCAACAACTCAGAGTAGAATCATCATCTAAATTGTGGGCTCAATGTGTCCAGTTACACAATGACATTCTCTTAGCTAAAGATACTACTGAAGCCTTTGAAAAAATGGTTTCACTACTTTCTGTTTTGCTTTCCATGCAGGGTGCTGTAGACATAAACAAGCTTTGTGAAGAAATGCTGGACAACAGGGCAACCTTACAAGCTATAGCCTCAGAGTTTAGTTCCCTTCCATCATATGCAGCTTTTGCTACTGCTCAAGAAGCTTATGAGCAGGCTGTTGCTAATGGTGATTCTGAAGTTGTTCTTAAAAAGTTGAAGAAGTCTTTGAATGTGGCTAAATCTGAATTTGACCGTGATGCAGCCATGCAACGTAAGTTGGAAAAGATGGCTGATCAAGCTATGACCCAAATGTATAAACAGGCTAGATCTGAGGACAAGAGGGCAAAAGTTACTAGTGCTATGCAGACAATGCTTTTCACTATGCTTAGAAAGTTGGATAATGATGCACTCAACAACATTATCAACAATGCAAGAGATGGTTGTGTTCCCTTGAACATAATACCTCTTACAACAGCAGCCAAACTAATGGTTGTCATACCAGACTATAACACATATAAAAATACGTGTGATGGTACAACATTTACTTATGCATCAGCATTGTGGGAAATCCAACAGGTTGTAGATGCAGATAGTAAAATTGTTCAACTTAGTGAAATTAGTATGGACAATTCACCTAATTTAGCATGGCCTCTTATTGTAACAGCTTTAAGGGCCAATTCTGCTGTCAAATTACAGAATAATGAGCTTAGTCCTGTTGCACTACGACAGATGTCTTGTGCTGCCGGTACTACACAAACTGCTTGCACTGATGACAATGCGTTAGCTTACTACAACACAACAAAGGGAGGTAGGTTTGTACTTGCACTGTTATCCGATTTACAGGATTTGAAATGGGCTAGATTCCCTAAGAGTGATGGAACTGGTACTATCTATACAGAACTGGAACCACCTTGTAGGTTTGTTACAGACACACCTAAAGGTCCTAAAGTGAAGTATTTATACTTTATTAAAGGATTAAACAACCTAAATAGAGGTATGGTACTTGGTAGTTTAGCTGCCACAGTACGTCTACAAGCTGGTAATGCAACAGAAGTGCCTGCCAATTCAACTGTATTATCTTTCTGTGCTTTTGCTGTAGATGCTGCTAAAGCTTACAAAGATTATCTAGCTAGTGGGGGACAACCAATCACTAATTGTGTTAAGATGTTGTGTACACACACTGGTACTGGTCAGGCAATAACAGTTACACCGGAAGCCAATATGGATCAAGAATCCTTTGGTGGTGCATCGTGTTGTCTGTACTGCCGTTGCCACATAGATCATCCAAATCCTAAAGGATTTTGTGACTTAAAAGGTAAGTATGTACAAATACCTACAACTTGTGCTAATGACCCTGTGGGTTTTACACTTAAAAACACAGTCTGTACCGTCTGCGGTATGTGGAAAGGTTATGGCTGTAGTTGTGATCAACTCCGCGAACCCATGCTTCAGTCAGCTGATGCACAATCGTTTTTAAACGGGTTTGCGGTGTAAGTGCAGCCCGTCTTACACCGTGCGGCACAGGCACTAGTACTGATGTCGTATACAGGGCTTTTGACATCTACAATGATAAAGTAGCTGGTTTTGCTAAATTCCTAAAAACTAATTGTTGTCGCTTCCAAGAAAAGGACGAAGATGACAATTTAATTGATTCTTACTTTGTAGTTAAGAGACACACTTTCTCTAACTACCAACATGAAGAAACAATTTATAATTTACTTAAGGATTGTCCAGCTGTTGCTAAACATGACTTCTTTAAGTTTAGAATAGACGGTGACATGGTACCACATATATCACGTCAACGTCTTACTAAATACACAATGGCAGACCTCGTCTATGCTTTAAGGCATTTTGATGAAGGTAATTGTGACACATTAAAAGAAATACTTGTCACATACAATTGTTGTGATGATGATTATTTCAATAAAAAGGACTGGTATGATTTTGTAGAAAACCCAGATATATTACGCGTATACGCCAACTTAGGTGAACGTGTACGCCAAGCTTTGTTAAAAACAGTACAATTCTGTGATGCCATGCGAAATGCTGGTATTGTTGGTGTACTGACATTAGATAATCAAGATCTCAATGGTAACTGGTATGATTTCGGTGATTTCATACAAACCACGCCAGGTAGTGGAGTTCCTGTTGTAGATTCTTATTATTCATTGTTAATGCCTATATTAACCTTGACCAGGGCTTTAACTGCAGAGTCACATGTTGACACTGACTTAACAAAGCCTTACATTAAGTGGGATTTGTTAAAATATGACTTCACGGAAGAGAGGTTAAAACTCTTTGACCGTTATTTTAAATATTGGGATCAGACATACCACCCAAATTGTGTTAACTGTTTGGATGACAGATGCATTCTGCATTGTGCAAACTTTAATGTTTTATTCTCTACAGTGTTCCCACTTACAAGTTTTGGACCACTAGTGAGAAAAATATTTGTTGATGGTGTTCCATTTGTAGTTTCAACTGGATACCACTTCAGAGAGCTAGGTGTTGTACATAATCAGGATGTAAACTTACATAGCTCTAGACTTAGTTTTAAGGAATTACTTGTGTATGCTGCTGACCCTGCTATGCACGCTGCTTCTGGTAATCTATTACTAGATAAACGCACTACGTGCTTTTCAGTAGCTGCACTTACTAACAATGTTGCTTTTCAAACTGTCAAACCCGGTAATTTTAACAAAGACTTCTATGACTTTGCTGTGTCTAAGGGTTTCTTTAAGGAAGGAAGTTCTGTTGAATTAAAACACTTCTTCTTTGCTCAGGATGGTAATGCTGCTATCAGCGATTATGACTACTATCGTTATAATCTACCAACAATGTGTGATATCAGACAACTACTATTTGTAGTTGAAGTTGTTGATAAGTACTTTGATTGTTACGATGGTGGCTGTATTAATGCTAACCAAGTCATCGTCAACAACCTAGACAAATCAGCTGGTTTTCCATTTAATAAATGGGGTAAGGCTAGACTTTATTATGATTCAATGAGTTATGAGGATCAAGATGCACTTTTCGCATATACAAAACGTAATGTCATCCCTACTATAACTCAAATGAATCTTAAGTATGCCATTAGTGCAAAGAATAGAGCTCGCACCGTAGCTGGTGTCTCTATCTGTAGTACTATGACCAATAGACAGTTTCATCAAAAATTATTGAAATCAATAGCCGCCACTAGAGGAGCTACTGTAGTAATTGGAACAAGCAAATTCTATGGTGGTTGGCACAACATGTTAAAAACTGTTTATAGTGATGTAGAAAACCCTCACCTTATGGGTTGGGATTATCCTAAATGTGATAGAGCCATGCCTAACATGCTTAGAATTATGGCCTCACTTGTTCTTGCTCGCAAACATACAACGTGTTGTAGCTTGTCACACCGTTTCTATAGATTAGCTAATGAGTGTGCTCAAGTATTGAGTGAAATGGTCATGTGTGGCGGTTCACTATATGTTAAACCAGGTGGAACCTCATCAGGAGATGCCACAACTGCTTATGCTAATAGTGTTTTTAACATTTGTCAAGCTGTCACGGCCAATGTTAATGCACTTTTATCTACTGATGGTAACAAAATTGCCGATAAGTATGTCCGCAATTTACAACACAGACTTTATGAGTGTCTCTATAGAAATAGAGATGTTGACACAGACTTTGTGAATGAGTTTTACGCATATTTGCGTAAACATTTCTCAATGATGATACTCTCTGACGATGCTGTTGTGTGTTTCAATAGCACTTATGCATCTCAAGGTCTAGTGGCTAGCATAAAGAACTTTAAGTCAGTTCTTTATTATCAAAACAATGTTTTTATGTCTGAAGCAAAATGTTGGACTGAGACTGACCTTACTAAAGGACCTCATGAATTTTGCTCTCAACATACAATGCTAGTTAAACAGGGTGATGATTATGTGTACCTTCCTTACCCAGATCCATCAAGAATCCTAGGGGCCGGCTGTTTTGTAGATGATATCGTAAAAACAGATGGTACACTTATGATTGAACGGTTCGTGTCTTTAGCTATAGATGCTTACCCACTTACTAAACATCCTAA TCAGGAGTATGCTGATGTCTTTCATTTGTACTTACAATACATAAGAAAGCTACATGATGAGTTAACAGGACACATGTTAGACATGTATTCTGTTATGCTTACTAATGATAACACTTCAAGGTATTGGGAACCTGAGTTTTATGAGGCTATGTACACACCGCATACAGTCTTACAGGCTGTTGGGGCTTGTGTTCTTTGCAATTCACAGACTTCATTAAGATGTGGTGCTTGCATACGTAGACCATTCTTATGTTGTAAATGCTGTTACGACCATGTCATATCAACATCACATAAATTAGTCTTGTCTGTTAATCCGTATGTTTGCAATGCTCCAGGTTGTGATGTCACAGATGTGACTCAACTTTACTTAGGAGGTATGAGCTATTATTGTAAATCACATAAACCACCCATTAGTTTTCCATTGTGTGCTAATGGACAAGTTTTTGGTTTATATAAAAATACATGTGTTGGTAGCGATAATGTTACTGACTTTAATGCAATTGCAACATGTGACTGGACAAATGCTGGTGATTACATTTTAGCTAACACCTGTACTGAAAGACTCAAGCTTTTTGCAGCAGAAACGCTCAAAGCTACTGAGGAGACATTTAAACTGTCTTATGGTATTGCTACTGTACGTGAAGTGCTGTCTGACAGAGAATTACATCTTTCATGGGAAGTTGGTAAACCTAGACCACCACTTAACCGAAATTATGTCTTTACTGGTTATCGTGTAACTAAAAACAGTAAAGTACAAATAGGAGAGTACACCTTTGAAAAAGGTGACTATGGTGATGCTGTTGTTTACCGAGGTACAACAACTTACAAATTAAATGTTGGTGATTATTTTGTGCTGACATCACATACAGTAATGCCATTAAGTGCACCTACACTAGTGCCACAAGAGCACTATGTTAGAATTACTGGCTTATACCCAACACTCAATATCTCAGATGAGTTTTCTAGCAATGTTGCAAATTATCAAAAGGTTGGTATGCAAAAGTATTCTACACTCCAGGGACCACCTGGTACTGGTAAGAGTCATTTTGCTATTGGCCTAGCTCTCTACTACCCTTCTGCTCGCATAGTGTATACAGCTTGCTCTCATGCCGCTGTTGATGCACTATGTGAGAAGGCATTAAAATATTTGCCTATAGATAAATGTAGTAGAATTATACCTGCACGTGCTCGTGTAGAGTGTTTTGATAAATTCAAAGTGAATTCAACATTAGAACAGTATGTCTTTTGTACTGTAAATGCATTGCCTGAGACGACAGCAGATATAGTTGTCTTTGATGAAATTTCAATGGCCACAAATTATGATTTGAGTGTTGTCAATGCCAGATTACGTGCTAAGCACTATGTGTACATTGGCGACCCTGCTCAATTACCTGCACCACGCACATTGCTAACTAAGGGCACACTAGAACCAGAATATTTCAATTCAGTGTGTAGACTTATGAAAACTATAGGTCCAGACATGTTCCTCGGAACTTGTCGGCGTTGTCCTGCTGAAATTGTTGACACTGTGAGTGCTTTGGTTTATGATAATAAGCTTAAAGCACATAAAGACAAATCAGCTCAATGCTTTAAAATGTTTTATAAGGGTGTTATCACGCATGATGTTTCATCTGCAATTAACAGGCCACAAATAGGCGTGGTAAGAGAATTCCTTACACGTAACCCTGCTTGGAGAAAAGCTGTCTTTATTTCACCTTATAATTCACAGAATGCTGTAGCCTCAAAGATTTTGGGACTACCAACTCAAACTGTTGATTCATCACAGGGCTCAGAATATGACTATGTCATATTCACTCAAACCACTGAAACAGCTCACTCTTGTAATGTAAACAGATTTAATGTTGCTATTACCAGAGCAAAAGTAGGCATACTTTGCATAATGTCTGATAGAGACCTTTATGACAAGTTGCAATTTACAAGTCTTGAAATTCCACGTAGGAATGTGGCAACTTTACAAGCTGAAAATGTAACAGGACTCTTTAAAGATTGTAGTAAGGTAATCACTGGATTACATCCTACACAGGCACCTACACACCTCAGTGTTGACACTAAATTCAAAACTGAAGGTTTATGTGTTGACATACCTGGCATACCTAAGGACATGACCTATAGAAGACTCATCTCTATGATGGGTTTTAAAATGAATTATCAAGTTAATGGTTACCCTAACATGTTTATCACCCGCGAAGAAGCTATAAGACATGTACGTGCATGGATTGGCTTCGATGTCGAGGGGTGTCATGCTACTAGAGAAGCTGTTGGTACCAATTTACCTTTACAGCTAGGTTTTTCTACAGGTGTTAACCTAGTTGCTGTACCTACAGGTTATGTTGATACACCTAATAATACAGATTTTTCCAGAGTTAGTGCTAAACCACCGCCTGGAGATCAATTTAAACACCTCATACCACTTATGTACAAAGGACTTCCTTGGAATGTAGTGCGTATAAAGATTGTACAAATGTTAAGTGACACACTTAAAAATCTCTCTGACAGAGTCGTATTTGTCTTATGGGCACATGGCTTTGAGTTGACATCTATGAAGTATTTTGTGAAAATAGGACCTGAGCGCACCTGTTGTCTATGTGATAGACGTGCCACATGCTTTTCCACTGCTTCAGACACTTATGCCTGTTGGCATCATTCTATTGGATTTGATTACGTCTATAATCCGTTTATGATTGATGTTCAACAATGGGGTTTTACAGGTAACCTACAAAGCAACCATGATCTGTATTGTCAAGTCCATGGTAATGCACATGTAGCTAGTTGTGATGCAATCATGACTAGGTGTCTAGCTGTCCACGAGTGCTTTGTTAAGCGTGTTGACTGGACTATTGAATATCCTATAATTGGTGATGAACTGAAGATTAATGCGGCTTGTAGAAAGGTTCAACACATGGTTGTTAAAGCTGCATTATTAGCAGACAAATTCCCAGTTCTTCACGACATTGGTAACCCTAAAGCTATTAAGTGTGTACCTCAAGCTGATGTAGAATGGAAGTTCTATGATGCACAGCCTTGTAGTGACAAAGCTTATAAAATAGAAGAATTATTCTATTCTTATGCCACACATTCTGACAAATTCACAGATGGTGTATGCCTATTTTGGAATTGCAATGTCGATAGATATCCTGCTAATTCCATTGTTTGTAGATTTGACACTAGAGTGCTATCTAACCTTAACTTGCCTGGTTGTGATGGTGGCAGTTTGTATGTAAATAAACATGCATTCCACACACCAGCTTTTGATAAAAGTGCTTTTGTTAATTTAAAACAATTACCATTTTTCTATTACTCTGACAGTCCATGTGAGTCTCATGGAAAACAAGTAGTGTCAGATATAGATTATGTACCACTAAAGTCTGCTACGTGTATAACACGTTGCAATTTAGGTGGTGCTGTCTGTAGACATCATGCTAATGAGTACAGATTGTATCTCGATGCTTATAACATGATGATCTCAGCTGGCTTTAGCTTGTGGGTTTACAAACAATTTGATACTTATAACCTCTGGAACACTTTTACAAGACTTCAGAGTTTAGAAAATGTGGCTTTTAATGTTGTAAATAAGGGACACTTTGATGGACAACAGGGTGAAGTACCAGTTTCTATCATTAATAACACTGTTTACACAAAAGTTGATGGTGTTGATGTAGAATTGTTTGAAAATAAAACAACATTACCTGTTAATGTAGCATTTGAGCTTTGGGCTAAGCGCAACATTAAACCAGTACCAGAGGTGAAAATACTCAATAATTTGGGTGTGGACATTGCTGCTAATACTGTGATCTGGGACTACAAAAGAGATGCTCCAGCACATATATCTACTATTGGTGTTTGTTCTATGACTGACATAGCCAAGAAACCAACTGAAACGATTTGTGCACCACTCACTGTCTTTTTTGATGGTAGAGTTGATGGTCAAGTAGACTTATTTAGAAATGCCCGTAATGGTGTTCTTATTACAGAAGGTAGTGTTAAAGGTTTACAACCATCTGTAGGTCCCAAACAAGCTAGTCTTAATGGAGTCACATTAATTGGAGAAGCCGTAAAAACACAGTTCAATTATTATAAGAAAGTTGATGGTGTTGTCCAACAATTACCTGAAACTTACTTTACTCAGAGTAGAAATTTACAAGAATTTAAACCCAGGAGTCAAATGGAAATTGATTTCTTAGAATTAGCTATGGATGAATTCATTGAACGGTATAAATTAGAAGGCTATGCCTTCGAACATATCGTTTATGGAGATTTTAGTCATAGTCAGTTAGGTGGTTTACATCTACTGATTGGACTAGCTAAACGTTTTAAGGAATCACCTTTTGAATTAGAAGATTTTATTCCTATGGACAGTACAGTTAAAAACTATTTCATAACAGATGCGCAAACAGGTTCATCTAAGTGTGTGTGTTCTGTTATTGATTTATTACTTGATGATTTTGTTGAAATAATAAAATCCCAAGATTTATCTGTAGTTTCTAAGGTTGTCAAAGTGACTATTGACTATACAGAAATTTCATTTATGCTTTGGTGTAAAGATGGCCATGTAGAAACATTTTACCCAAAATTACAATCTAGTCAAGCGTGGCAACCGGGTGTTGCTATGCCTAATCTTTACAAAATGCAAAGAATGCTATTAGAAAAGTGTGACCTTCAAAATTATGGTGATAGTGCAACATTACCTAAAGGCATAATGATGAATGTCGCAAAATATACTCAACTGTGTCAATATTTAAACACATTAACATTAGCTGTACCCTATAATATGAGAGTTATACATTTTGGTGCTGGTTCTGATAAAGGAGTTGCACCAGGTACAGCTGTTTTAAGACAGTGGTTGCCTACGGGTACGCTGCTTGTCGATTCAGATCTTAATGACTTTGTCTCTGATGCAGATTCAACTTTGATTGGTGATTGTGCAACTGTACATACAGCTAATAAATGGGATCTCATTATTAGTGATATGTACGACCCTAAGACTAAAAATGTTACAAAAGAAAATGACTCTAAAGAGGGTTTTTTCACTTACATTTGTGGGTTTATACAACAAAAGCTAGCTCTTGGAGGTTCCGTGGCTATAAAGATAACAGAACATTCTTGGAATGCTGATCTTTATAAGCTCATGGGACACTTCGCATGGTGGACAGCCTTTGTTACTAATGTGAATGCGTCATCATCTGAAGCATTTTTAATTGGATGTAATTATCTTGGCAAACCACGCGAACAAATAGATGGTTATGTCATGCATGCAAATTACATATTTTGGAGGAATACAAATCCAATTCAGTTGTCTTCCTATTCTTTATTTGACATGAGTAAATTTCCCCTTAAATTAAGGGGTACTGCTGTTATGTCTTTAAAAGAAGGTCAAATCAATGATATGATTTTATCTCTTCTTAGTAAAGGTAGACTTATAATTAGAGAAAACAACAGAGTTGTTATTTCTAGTGATGTTCTTGTTAACAACTAAACGAACAATGTTTGTTTTTCTTGTTTTATTGCCACTATAAGTAAGTCTCTAGTCAGTGTGTTAATTTTACAACCAGAACTCAATTACCCCCTGCATACACTAATTCTTTCACACGTGGTGTTTATTACCCTGACAAAGTTTTCAGATCCTCAGTTTTACATTCAACTCAGGACTTGTTCTTACCTTTCTTTTCCAATGTTACTTGGTTCCATGCTATACATGTCTCTGGGACCAATGGTACTAAGAGGTTTGCTAACCCTGTCCTACCATTTAATGATGGTGTTTATTTTGCTTCCACTGAGAAGTCTAACATAATAAGAGGCTGGATTTTTGGTACTACTTTAGATTCGAAGACCCAGTCCCTACTTATTGTTAATAACGCTACTAATGTTGTTATTAAAGTCTGTGAATTTCAATTTTGTAATGATCCATTTTTGGGTGTTTATTACCACAAAAACAACAAAAGTTGGATGGAAAGTGAGTTCAGAGTTTATTCTAGTGCGAATAATTGCACTTTTGAATATGTCTCTCAGCCTTTTCTTATGGACCTTGAAGGAAAACAGGGTAATTTCAAAAATCTTAGGGAATTTGTGTTTAAGAATATTGATGGTTATTTTAAAATATATTCTAAGCACACGCCTATTAATTTAGTGCGTGGTCTCCCTCAGGGTTTTTCGGCTTTAGAACCATTGGTAGATTTGCCAATAGGTATTAACATCACTAGGTTTCAAACTTTACATAGAAGTTATTTGACTCCTGGTGATTCTTCTTCAGGTTGGACAGCTGGTGCTGCAGCTTATTATGTGGGTTATCTTCAACCTAGGACTTTTCTATTAAAATATAATGAAAATGGAACCATTACAGATGCTGTAGACTGTGCACTTGACCCTCTCTCAGAAACAAAGTGTACGTTGAAATCCTTCACTGTAGAAAAAGGAATCTATCAAACTTCTAACTTTAGAGTCCAACCAACAGAATCTATTGTTAGATTTCCTAATATTACAAACTTGTGCCCTTTTGGTGAAGTTTTTAACGCCACCAGATTTGCATCTGTTTATGCTTGGAACAGGAAGAGAATCAGCAACTGTGTTGCTGATTATTCTGTCCTATATAATTCCGCATCATTTTCCACTTTTAAGTGTTATGGAGTGTCTCCTACTAAATTAAATGATCTCTGCTTTACTAATGTCTATGCAGATTCATTTGTAATTAGAGGTGATGAAGTCAGACAAATCGCTCCAGGGCAAACTGGAAATATTGCTGATTATAATTATAAATTACCAGATGATTTTACAGGCTGCGTTATAGCTTGGAATTCTAACAATCTTGATTCTAAGGTTGGTGGTAATTATAATTACCTGTATAGATTGTTTAGGAAGTCTAATCTCAAACCTTTTGAGAGAGATATTTCAACTGAAATCTATCAGGCCGGTAGCACACCTTGTAATGGTGTTAAAGGTTTTAATTGTTACTTTCCTTTACAATCATATGGTTTCCAACCCACTTATGGTGTTGGTTACCAACCATACAGAGTAGTAGTACTTTCTTTTGAACTTCTACATGCACCAGCAACTGTTTGTGGACCTAAAAAGTCTACTAATTTGGTTAAAAACAAATGTGTCAATTTCAACTTCAATGGTTTAACAGGCACAGGTGTTCTTACTGAGTCTAACAAAAAGTTTCTGCCTTTCCAACAATTTGGCAGAGACATTGCTGACACTACTGATGCTGTCCGTGATCCACAGACACTTGAGATTCTTGACATTACACCATGTTCTTTTGGTGGTGTCAGTGTTATAACACCAGGAACAAATACTTCTAACCAGGTTGCTGTTCTTTATCAGGGTGTTAACTGCACAGAAGTCCCTGTTGCTATTCATGCAGATCAACTTACTCCTACTTGGCGTGTTTATTCTACAGGTTCTAATGTTTTTCAAACACGTGCAGGCTGTTTAATAGGGGCTGAACATGTCAACAACTCATATGAGTGTGACATACCCATTGGTGCAGGTATATGCGCTAGTTATCAGACTCAGACTAATTCTCGTAGTGTAGCTAGTCAATCCATCATTGCCTACACTATGTCACTTGGTGTAGAAAATTCAGTTGCTTACTCTAATAACTCTATTGCCATACCCACAAATTTTACTATTAGTGTTACCACAGAAATTCTACCAGTGTCTATGACCAAGACATCAGTAGATTGTACAATGTACATTTGTGGTGATTCAACTGAATGCAGCAATCTTTTGTTGCAATATGGCAGTTTTTGTACACAATTAAACCGTGCTTTAACTGGAATAGCTGTTGAACAAGACAAAAACACCCAAGAAGTTTTTGCACAAGTCAAACAAATTTACAAAACACCACCAATTAAAGATTTTGGTGGTTTTAATTTTTCACAAATATTACCAGATCCATCAAAACCAAGCAAGAGGTCATTTATTGAAGATCTACTTTTCAACAAAGTGACACTTGCAGATGCTGGCTTCATCAAACAATATGGTGATTGCCTTGGTGATATTGCTGCTAGAGACCTCATTTGTGCACAAAAGTTTAACGGCCTTACTGTTTTGCCACCTTTGCTCACAGATGAAATGATTGCTCAATACACTTCTGCACTGTTAGCGGGTACAATCACTTCTGGTTGGACCTTTGGTGCAGGTGCTGCATTACAAATACCATTTGCTATGCAAATGGCTTATAGGTTTAATGGTATTGGAGTTACACAGAATGTTCTCTATGAGAACCAAAAATTGATTGCCAACCAATTTAATAGTGCTATTGGCAAAATTCAAGACTCACTTTCTTC CACAGCAAGTGCACTTGGAAAACTTCAAGATGTGGTCAACCAAAATGCACAAGCTTTAAACACGCTTGTTAAACAACTTAGCTCCAATTTTGGTGCAATTTCAAGTGTTTTAAATGATATCCTTTCACGTCTTGACAAAGTTGAGGCTGAAGTGCAAATTGATAGGTTGATCACAGGCAGACTTCAAAGTTTGCAGACATATGTGACTCAACAATTAATTAGAGCTGCAGAAATCAGAGCTTCTGCTAATCTTGCTGCTACTAAAATGTCAGAGTGTGTACTTGGACAATCAAAAAGAGTTGATTTTTGTGGAAAGGGCTATCATCTTATGTCCTTCCCTCAGTCAGCACCTCATGGTGTAGTCTTCTTGCATGTGACTTATGTCCCTGCACAAGAAAAGAACTTCACAACTGCTCCTGCCATTTGTCATGATGGAAAAGCACACTTTCCTCGTGAAGGTGTCTTTGTTTCAAATGGCACACACTGGTTTGTAACACAAAGGAATTTTTATGAACCACAAATCATTACTACAGACAACACATTTGTGTCTGGTAACTGTGATGTTGTAATAGGAATTGTCAACAACACAGTTTATGATCCTTTGCAACCTGAATTAGACTCATTCAAGGAGGAGTTAGATAAATATTTTAAGAATCATACATCACCAGATGTTGATTTAGGTGACATCTCTGGCATTAATGCTTCAGTTGTAAACATTCAAAAAGAAATTGACCGCCTCAATGAGGTTGCCAAGAATTTAAATGAATCTCTCATCGATCTCCAAGAACTTGGAAAGTATGAGCAGTATATAAAATGGCCATGGTACATTTGGCTAGGTTTTATAGCTGGCTTGATTGCCATAGTAATGGTGACAATTATGCTTTGCTGTATGACCAGTTGCTGTAGTTGTCTCAAGGGCTGTTGTTCTTGTGGATCCTGCTGCAAATTTGATGAAGACGACTCTGAGCCAGTGCTCAAAGGAGTCAAATTACATTACACATAAACGAACTTATGGATTTGTTTATGAGAATCTTCACAATTGGAACTGTAACTTTGAAGCAAGGTGAAATCAAGGATGCTACTCCTTCAGATTTTGTTCGCGCTACTGCAACGATACCGATACAAGCCTCACTCCCTTTCGGATGGCTTATTGTTGGCGTTGCACTTCTTGCTGTTTTTCATAGCGCTTCCAAAATCATAACCCTCAAAAAGAGATGGCAACTAGCACTCTCCAAGGGTGTTCACTTTGTTTGCAACTTGCTGTTGTTGTTTGTAACAGTTTACTCACACCTTTTGCTCGTTGCTGCTGGCCTTGAAGCCCCTTTTCTCTATCTTTATGCTTTAGTCTACTTCTTGCAGAGTATAAACTTTGTAAGAATAATAATGAGGCTTTGGCTTTGCTGGAAATGCCGTTCCAAAAACCCATTACTTTATGATGCCAACTATTTTCTTTGCTGGCATACTAATTGTTACGACTATTGTATACCTTACAATAGTGTAACTTCTTCAATTGTCATTACTTTATGTGATGGCACAACAAGTCCTATTTCTGAACATGACTACCAGATTGGTGGTTATACTGAAAAATGGGAATCTGGAGTAAAAGACTGTGTTGTATTACACAGTTACTTCACTTCAGACTATTACCAGCTGTACTCAACTCAATTGAGTACAGACACTGGTGTTGAACATGTTACCTTCTTCATCTACAATAAAATTGTTGATGAGCCTGAAGAACATGTCCAAATTCACACAATCGACGGTTCATCCGGAGTTGTTAATCCAGTAATGGAACCAATTTATGATGAACCGACGACGACTACTAGCGTGCCTTTGTAAGCACAAGCTGATGAGTACGAACTTATGAAAATTATTCTTTTCTTGGCACTGATAACACTCATGGTGAGCAAGGGCGAGGAGCTGTTCACCGGGGTGGTGCCCATCCTGGTCGAGCTGGACGGCGACGTAAACGGCCACAAGTTCAGCGTGTCCGGCGAGGGCGAGGGCGATGCCACCTACGGCAAGCTGACCCTGAAGTTCATCTGCACCACCGGCAAGCTGCCCGTGCCCTGGCCCACCCTCGTGACCACCCTGACCTACGGCGTGCAGTGCTTCAGCCGCTACCCCGACCACATGAAGCAGCACGACTTCTTCAAGTCCGCCATGCCCGAAGGCTACGTCCAGGAGCGCACCATCTTCTTCAAGGACGACGGCAACTACAAGACCCGCGCCGAGGTGAAGTTCGAGGGCGACACCCTGGTGAACCGCATCGAGCTGAAGGGCATCGACTTCAAGGAGGACGGCAACATCCTGGGGCACAAGCTGGAGTACAACTACAACAGCCACAACGTCTATATCATGGCCGACAAGCAGAAGAACGGCATCAAGGTGAACTTCAAGATCCGCCACAACATCGAGGACGGCAGCGTGCAGCTCGCCGACCACTACCAGCAGAACACGCCGATTGGTGATGGTCCGGTGCTGCTGCCCGACAACCACTACCTGAGCACCCAGTCCGCCCTGAGCAAAGACCCCAACGAGAAGCGCGATCACATGGTCCTGCTGGAGTTCGTGACCGCCGCCGGGATCACTCTCGGCATGGACGAGCTGTACAAGGTCTTCACACTCGAAGATTTCGTTGGGGACTGGCGACAGACAGCCGGCTACAACCTGGACCAAGTCCTTGAACAGGGAGGTGTGTCCAGTTTGTTTCAGAATCTCGGGGTGTCCGTAACTCCGATCCAAAGGATTGTCCTGAGCGGTGAAAATGGGCTGAAGATCGACATCCATGTCATCATCCCGTATGAAGGTCTGAGCGGCGACCAAATGGGCCAGATCGAAAAAATTTTTAAGGTGGTGTACCCTGTGGATGATCATCACTTTAAGGTGATCCTGCACTATGGCACACTGGTAATCGACGGGGTTACGCCGAACATGATCGACTATTTCGGACGGCCGTATGAAGGCATCGCCGTGTTCGACGGCAAAAAGATCACTGTAACAGGGACCCTGTGGAACGGCAACAAAATTATCGACGAGCGCCTGATCAACCCCGACGGCTCCCTGCTGTTCCGAGTAACCATCAACGGAGTGACCGGCTGGCGGCTGTGCGAACGCATTCTGGCGTAAGTGACAACAGATGTTTCATCTCGTTGACTTTCAGGTTACTATAGCAGAGATATTACTAATTATTATGAGGACTTTTAAAGTTTCCATTTGGAATCTTGATTACATCATAAACCTCATAATTAAAAATTTATCTAAGTCACTAACTGAGAATAAATATTCTCAATTAGATGAAGAGCAACCAATGGAGATTGATTAAACGAACATGAAAATTATTCTTTTCTTGGCACTGATAACACTCGCTACTTGTGAGCTTTATCACTACCAAGAGTGTGTTAGAGGTACAACAGTACTTTTAAAAGAACCTTGCTCTTCTGGAACATACGAGGGCAATTCACCATTTCATCCTCTAGCTGATAACAAATTTGCACTGACTTGCTTTAGCACTCAATTTGCTTTTGCTTGTCCTGACGGCGTAAAACACGTCTATCAGTTACGTGCCAGATCAGTTTCACCTAAACTGTTCATCAGACAAGAGGAAGTTCAAGAACTTTACTCTCCAATTTTTCTTATTGTTGCGGCAATAGTGTTTATAACACTTTGCTTCACACTCAAAAGAAAGACAGAATGATTGAACTTTCATTAATTGACTTCTATTTGTGCTTTTTAGCCTTTCTGCTATTCCTTGTTTTAATTATGCTTATTATCTTTTGGTTCTCACTTGAACTGCAAGATCATAATGAAACTTGTCACGCCTAAACGAACATGAAATTTCTTGTTTTCTTAGGAATCATCACAACTGTAGCTGCATTTCACCAAGAATGTAGTTTACAGTCATGTACTCAACATCAACCATATGTAGTTGATGACCCGTGTCCTATTCACTTCTATTCTAAATGGTATATTAGAGTAGGAGCTAGAAAATCAGCACCTTTAATTGAATTGTGCGTGGATGAGGCTGGTTCTAAATCACCCATTCAGTACATCGATATCGGTAATTATACAGTTTCCTGTTTACCTTTTACAATTAATTGCCAGGAACCTAAATTGGGTAGTCTTGTAGTGCGTTGTTCGTTCTATGAAGACTTTTTAGAGTATCATGACGTTCGTGTTGTTTTAGATTTTATCTAAACGAACAAACTAAAATGTCTGATAATGGACCCCAAAATCAGCGAAATGCACCCCGCATTACGTTTGGTGGACCCTCAGATTCAACTGGCAGTAACCAGAATGGAGAACGCAGTGGGGCGCGATCAAAACAACGTCGGCCCCAAGGTTTACCCAATAATACTGCGTCTTGGTTCACCGCTCTCACTCAACATGGCAAGGAAGACCTTAAATTCCCTCGAGGACAAGGCGTTCCAATTAACACCAATAGCAGTCCAGATGACCAAATTGGCTACTACCGAAGAGCTACCAGACGAATTCGTGGTGGTGACGGTAAAATGAAAGATCTCAGTCCAAGATGGTATTTCTACTACCTAGGAACTGGGCCAGAAGCTGGACTTCCCTATGGTGCTAACAAAGACGGCATCATATGGGTTGCAACTGAGGGAGCCTTGAATACACCAAAAGATCACATTGGCACCCGCAATCCTGCTAACAATGCTGCAATCGTGCTACAACTTCCTCAAGGAACAACATTGCCAAAAGGCTTCTACGCAGAAGGGAGCAGAGGCGGCAGTCAAGCCTCTTCTCGTTCCTCATCACGTAGTCGCAACAGTTCAAGAAATTCAACTCCAGGCAGCAGTAGGGGAATTTCTCCTGCTAGAATGGCTGGCAATGGCGGTGATGCTGCTCTTGCTTTGCTGCTGCTTGACAGATTGAACCAGCTTGAGAGCAAAATGTCTGGTAAAGGCCAACAACAACAAGGCCAAACTGTCACTAAGAAATCTGCTGCTGAGGCTTCTAAGAAGCCTCGGCAAAAACGTACTGCCACTAAAGCATACAATGTAACACAAGCTTTCGGCAGACGTGGTCCAGAACAAACCCAAGGAAATTTTGGGGACCAGGAACTAATCAGACAAGGAACTGATTACAAACATTGGCCGCAAATTGCACAATTTGCCCCCAGCGCTTCAGCGTTCTTCGGAATGTCGCGCATTGGCATGGAAGTCACACCTTCGGGAACGTGGTTGACCTACACAGGTGCCATCAAATTGGATGACAAAGATCCAAATTTCAAAGATCAAGTCATTTTGCTGAATAAGCATATTGACGCATACAAAACATTCCCACCAACAGAGCCTAAAAAGGACAAAAAGAAGAAGGCTGATGAAACTCAAGCCTTACCGCAGAGACAGAAGAAACAGCAAACTGTGACTCTTCTTCCTGCTGCAGATTTGGATGATTTCTCCAAACAATTGCAACAATCCATGAGCAGTGCTGACTCAACTCAGGCCTAAACTCATGCAGACCACACAAGGCAGATGGGCTATATAAACGTTTTCGCTTTTCCGTTTACGATATATAGTCTACTCTTGTGCAGAATGAATTCTCGTAACTACATAGCACAAGTAGATGTAGTTAACTTTAATCTCACATAGCAATCTTTAATCAGTGTGTAACATTAGGGAGGACTTGAAAGAGCCACCACATTTTCACCGAGGCCACGCGGAGTACGATCGAGTGTACAGTGAACAATGCTAGGGAGAGCTGCCTATATGGAAGAGCCCTAATGTGTAAAATTAATTTTAGTAGTGCTATCCCCATGTGATTTTAATAGCTTCTT AGGAGAATGACAA

